# VELCRO-IP RNA-seq explores ribosome expansion segment function in translation genome-wide

**DOI:** 10.1101/2020.07.01.179515

**Authors:** Kathrin Leppek, Gun Woo Byeon, Kotaro Fujii, Maria Barna

**Affiliations:** Department of Developmental Biology, Stanford University, Stanford, California 94305, USA; Department of Genetics, Stanford University, Stanford, California 94305, USA

**Keywords:** mRNA translation, ribosome, internal initiation, internal ribosome entry site, stem-loop, RNA structure, yeast, ribosomal RNA, ribosome engineering, expansion segment, ES9S, RNA sequencing, regional RNA fragment mapping, mouse embryo

## Abstract

Roles for ribosomal RNA (rRNA) in gene regulation remain largely unexplored. With hundreds of rDNA units scattered across multiple chromosomal loci, it is not possible to genetically modify rRNA in mammalian cells, hindering understanding of ribosome function. Emerging evidence suggests that expansion segments (ESs), tentacle-like rRNA extensions that vary in sequence and size across eukaryotic evolution, may provide platforms for the binding of proteins and mRNAs. Here, we develop VELCRO-IP RNA-seq: a versatile methodology to generate species-adapted ESs and map specific mRNA regions across the transcriptome that preferentially associate with ESs. By applying VELCRO-IP RNA-seq to a mammalian ES, ES9S, we identified a large array of mRNAs that are selectively recruited to ribosomes via an ES. We further characterize a set of specific 5’ UTRs that facilitate cap-independent translation through ES9S-mediated ribosome recruitment. These data provide a novel technology for studying the enigmatic ESs of the ribosome in gene-specific translation.

## INTRODUCTION

The ribosome is life’s most ancient molecular machine with an RNA structural core that is universally shared across all species. However, a dramatic increase in its size has occurred during eukaryotic evolution. For example, the human ribosome is 1MDa larger than the yeast ribosome which in turn is another 1MDa larger than the bacterial ribosome. This is due in large part to the insertions of blocks of sequences that are called expansion segments (ESs) as they “expand” the eukaryotic ribosomal RNA (rRNA) (Gerbi, 1996). ESs are located in rRNA regions of lower primary sequence conservation which suggests that they are neutral mutations at non-essential domains of the ribosome that do not interfere with essential rRNA function in translation across all kingdoms (Gerbi, 1986). However, it has not been determined whether ESs are generally dispensable or instead, have more specialized cellular functions. In particular, it remains largely unknown whether there are species-specific roles of ESs in regulation of translation. Although ESs are generally found at the same relative location in the rRNAs of different eukaryotes, they can exhibit a striking degree of variability in their length and sequence both within and among different species, including different tissue types (Kuo et al., 1996; Leffers and Andersen, 1993; Parks et al., 2018). For example, although yeast have a similar number of ESs as humans, ESs vary dramatically in sequence and size across eukaryotic evolution. The longest of ESs are more than 700 nt in *Homo sapiens* (*H. sapiens*) and resemble flexible “tentacles” that extend from the ribosome (Anger et al., 2013; Armache et al., 2010; Gerbi, 1996). The biological impact of ES variation on translation could be a critical facet of our understanding of the evolution of gene regulation and organismal development. For the last several decades however, directly addressing this question has been limited by the lack of a robust system to manipulate and investigate rRNA at the genetic and molecular level.

The major challenge in the investigation of ESs lies in that rRNAs are transcribed from ribosomal DNA (rDNA) loci that consist of hundreds of tandemly repeated units. rDNA copy number varies between eukaryotic species, ranging from a few hundred in most metazoans up to thousands of copies, for example, in wheat (Appels et al., 1980). Thus, for most higher eukaryotic species, it has not been possible to experimentally manipulate rRNA to identify mRNAs or proteins that interact with a specific ES within the context of the assembled ribosome. Here, we report the development of VELCRO (<u>v</u>ariable <u>e</u>xpansion <u>s</u>egment-ligand <u>c</u>himeric <u>r</u>ib<u>o</u>some), a versatile methodology to generate chimeric ribosomes in which the species-specific ES under investigation replaces its native counterpart in the yeast ribosome. We further show how such chimeric ribosomes can be coupled with a biochemical pulldown approach (VELCRO-IP) and RNA sequencing (VELCRO-IP RNA-seq) in a modular manner to interrogate rRNA-mRNA interactions genome-wide.

We have previously established the functional and mechanistic framework for the direct interaction of the mammalian ES9S of the 18S rRNA with a specific 5’ UTR RNA element, the short P4 stem-loop of the *Hoxa9* IRES-like element, for cap-independent translation initiation of the downstream mRNA coding region (Leppek et al., 2020). Through ES9S-mediated ribosome recruitment by *Hoxa9* P4, this RNA stem-loop acts as a translational enhancer. The functional importance of the *Hoxa9* P4-ES9S interaction for mammalian translation regulation suggests that additional mRNAs may potentially be regulated via ES9S-mediated recruitment. We thus chose ES9S as the proof-of-principle ES variant of interest to develop the presented technology. This provided us with both a confirmed ES-mRNA binding event necessary to verify our method and a key biological question that remained unanswered. By applying our VELCRO-IP RNA-seq technology to mammalian ES9S, we discover the unexpected function of ES9S in gene regulation, where transcriptome-wide binding of specific mRNAs to ES9S through their 5’ UTRs enable cap-independent translation of the mRNA in a species-specific manner. Our results highlight the role of the evolution of the ribosome ESs in guiding species- and gene-specific translation and provide a novel technology broadly applicable to investigations of enigmatic variations in rRNA.

## DESIGN

We set out to explore a potential broader function of the ribosome ESs to mediate, in a genome-wide fashion, species- and sequence-specific interaction between ribosomes and mRNAs. rRNA-mRNA interaction is a classic paradigm for translation initiation in prokaryotes where the Shine-Dalgarno leader sequence in mRNAs base-pairs with the 16S rRNA 3’ end to designate translation start sites (Shine and Dalgarno, 1974; Steitz and Jakes, 1975). But beyond a few individual mRNAs, no clear evidence for the transcriptome-wide use of such mechanisms have been demonstrated for any eukaryotic species. To obtain direct and functionally relevant biochemical evidence of rRNA-mRNA interactions genome-wide, the rRNA must be presented and probed in its native functional structure and in context of the ribosome. While eukaryotic ribosomes can be purified from cellular extracts, it has remained a challenge to obtain ribosomes with experimentally manipulated rRNA components in a sequence-specific manner. This lack of comprehensive methods to study ES function, or frankly any rRNA functions beyond peptide bond formation, is because genetic manipulation of ribosomal DNA (rDNA) regions has not been possible for most higher eukaryotes due to the repetitive nature of hundreds of rDNA units spread across multiple chromosomal loci in metazoans (Romanova et al., 2006). Thus, a method that overcomes these limitations is required to pursue the question of specific mRNA recruitment to the ribosome via an ES and to enable broader inquiries into the function of ribosome ESs in general, across species-, tissue- or individual-specific rDNA variants.

Baker’s yeast, *Saccharomyces cerevisiae* (*S. cerevisiae*), while still possessing a repetitive tandem array of rDNA units, contains a single rDNA locus in its genome. This locus has previously been deleted in its entirety and can be complemented with an exogenous rDNA expressing plasmid that enables genetic manipulation of ribosomes in yeasts (Nemoto et al., 2010; Wai et al., 2000). This led us to envision a strategy in which the variable ES of interest could replace the native ES sequence of the yeast rRNA through the rDNA complementation approach. In our development of the VELCRO method, we first examine the diversity in sequence and structure of ES9S, our paradigm example, across different species and engineer the chimeric ribosomes by “humanizing” yeast 18S rRNA exclusively in the distal part of ES9S (**Figure 1, S1**) (Leppek et al., 2020). We introduce an endogenous Flag tag to enable affinity purification of chimeric ribosomes, and we verify that the chimeric, Flag tagged ribosomes are incorporated into translating polysomes (Figure S2). We establish a pulldown method via the Flag tag for selectively purifying the interactors from the input pool of fragmented mouse embryo mRNAs (**Figure 2, 3**). We then utilize high throughput RNA sequencing to identify regions of the embryonic mRNAs that specifically interact with the humanized ES in a quantitative and genome-wide fashion (**Figure 4**). In our design of VELCRO-IP RNA-seq, we detect for each mRNA region how well it interacts with ribosomes containing human ES relative to its level of interaction with ribosomes containing the corresponding yeast ES to control for any potential confounding variables. Using replicate data, we calculate empirical estimates of statistical confidence that demonstrate high signal-to-noise ratio of our method and classify specifically and highly enriched mRNA regions that preferentially associate with the human ES (**Figure 4, 5**). Importantly, we also provide a reverse pulldown approach that allows orthogonal validation of the discovered mRNA-ES interactions using the same yeast strains used for VELCRO-IP (**Figure 6**). Together, our VELCRO-IP RNA-seq offers a versatile, modular, and rigorous methodology to investigate variations of the ribosome ESs.

**Figure 1.**
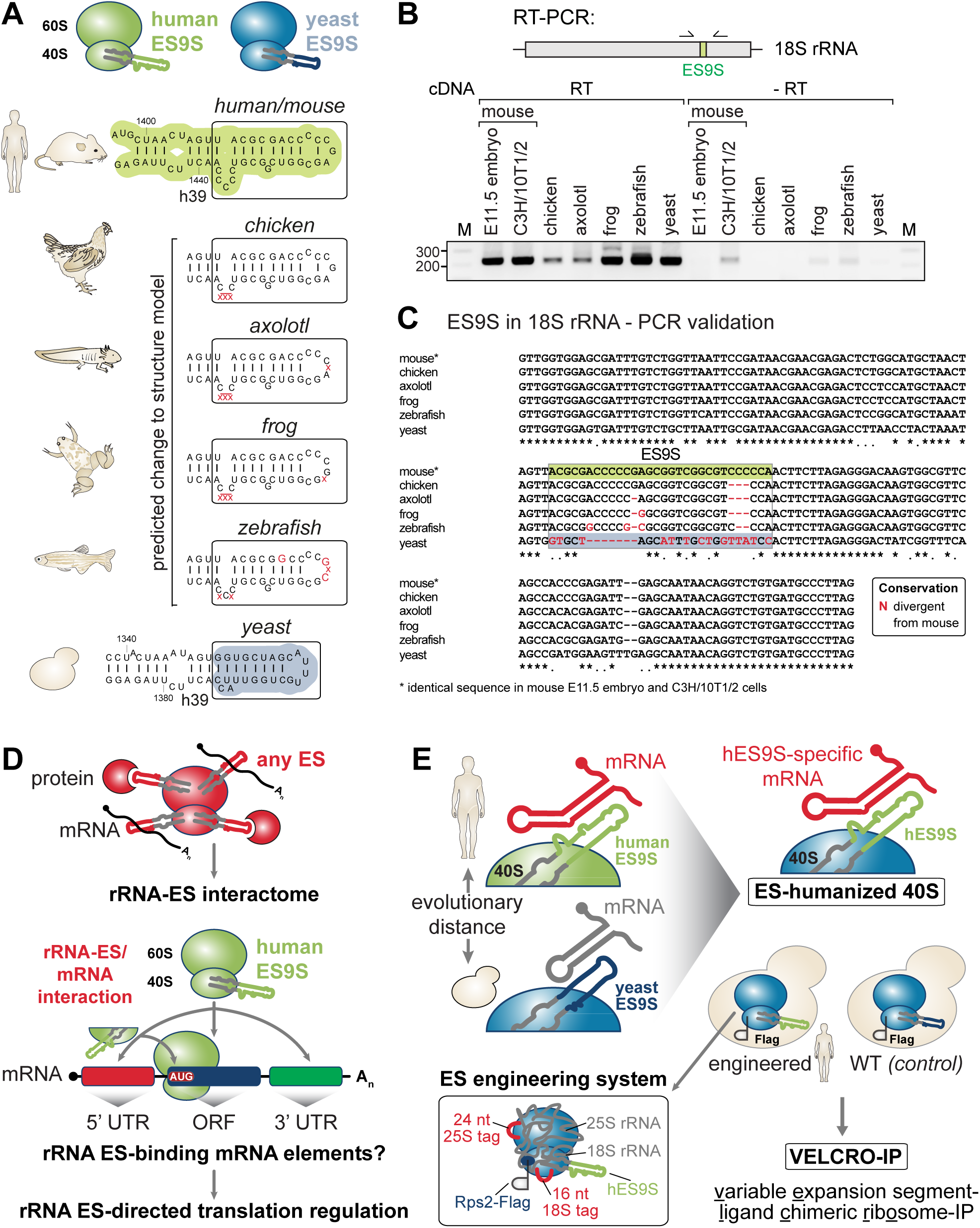
Confirmation of interspecies sequence variation of ES9S rRNA region. (A) Secondary structure models of the human (*H. sapiens*) and baker’s yeast (*S. cerevisiae*) 18S rRNA region containing ES9S, highlighted in green and blue, respectively. Predicted structural changes in the ES9S region of 18S rRNA with regards to species-specific variation in sequence. Sequence changes are annotated and their possible effect on the ES9S structure in red that are divergent from the identical human and mouse ES9S. Secondary structure models of ES9S of different species were predicted using Vienna RNAfold (http://rna.tbi.univie.ac.at) and visualized using VARNA (http://varna.lri.fr) with default settings. See also **Figure S1**. (B) Schematic of the RT-PCR analysis of the ES9S region in 18S rRNA using cDNA generated from total RNA derived from six different species (E11.5, stage E11.5 FVB mouse embryo; chicken, *Gallus gallus*; axolotl, *Ambystoma mexicanum*; frog, X. l., *Xenopus laevis*; zebrafish, *Danio rerio*; yeast, S. c., *Saccharomyces cerevisiae*) and primers specific for the 18S rRNA region containing ES9S in the center (see **Table S3**). (C) Multiple-species sequence alignment of the 18S rRNA region including ES9S indicating ES9S as a variable species-specific sequence embedded in highly conserved 18S rRNA sequence. Sequencing of the single PCR product with the outer primers from (B) confirms the species-specific ES9S sequence as annotated. For six different species we verified annotated 18S rRNA sequences by RT-PCR spanning the ES9S region. Nts divergent from mouse ES9S are highlighted in red. (D) Concept of exploring extended rRNA ES interactions on the ribosome with components such as mRNAs or proteins. With a focus on ES9S on the 40S ribosomal subunit we are investigating, as a paradigm example, ES9S-mRNA interactions via the ribosome with positional resolution in order to identify and map ES9S-binding mRNA element. ES-mRNA interactions may give insights into unexplored ES-directed translation regulation. (E) Schematic of our approach to investigate ES-mediated translation regulation through mRNA-interactions. The evolutionary distance between human and yeast ribosomes can be harnessed by “humanizing” yeast 18S rRNA exclusively in the exemplary ES9S region. By generating Flag-tagged humanized ribosome-strains alongside tagged WT control yeast strains we envision an ES engineering system that contains rRNA and protein tags and allows the manipulation of an ES of choice. The technology presented is termed VELCRO-IP: <u>v</u>ariable <u>e</u>xpansion segment-ligand <u>c</u>himeric <u>r</u>ib<u>o</u>some-IP.

**Figure 2.**
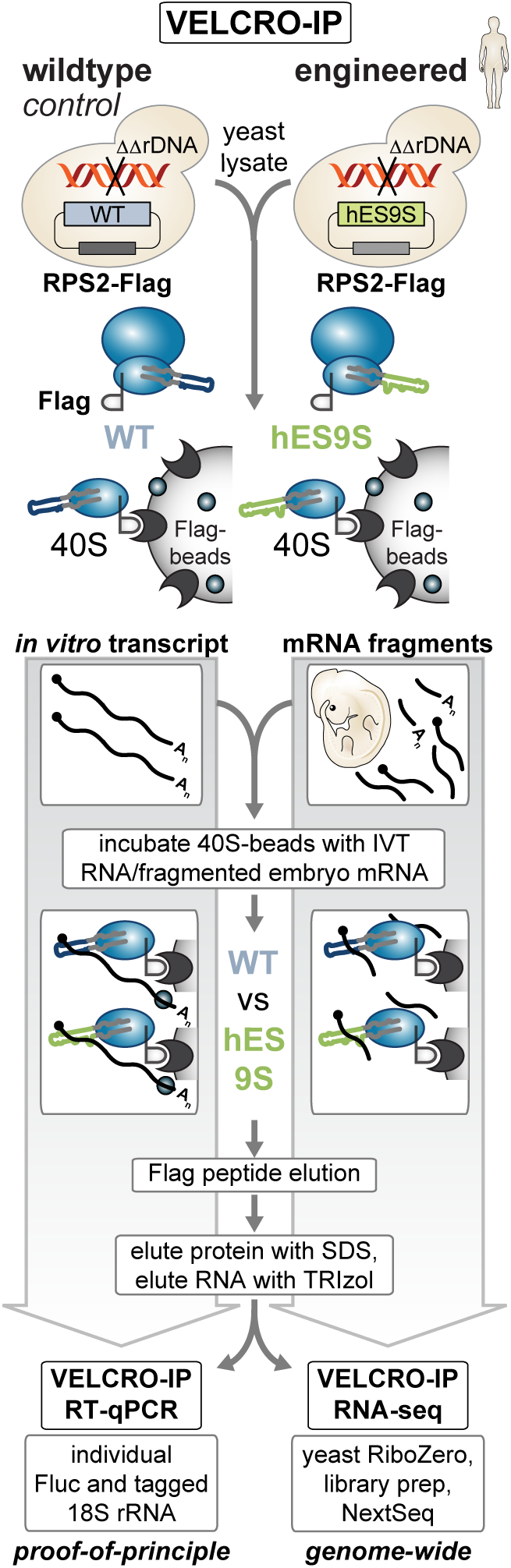
Development of VELCRO-IP RNA-seq to identify global ES-mRNA interactions. Schematic representation of the VELCRO-IP approach. 40S ribosomes are tagged by endogenously Flag-tagging Rps2 at the C-terminus and by generating WT or hES9S rDNA containing Rps2-Flag yeast strains. Lysates are generated by cryo-milling, and 40S ribosomal subunits from each strain are coupled to Flag agarose beads and washed. For a proof-of-principle VELCRO-IP RT-qPCR experiment, *in vitro* transcripts (IVT) described in Figure 3 are incubated with washed ribosome beads. Upon 3xFlag peptide-elution of 40S-RNA complexes, total RNA is eluted with TRIzol, and IVT enrichment is determined by RT-qPCR using primers specific for Fluc and the 18S rRNA tag to normalize for 40S-IP efficiency. For the genome-wide VELCRO-IP RNA-seq experiment, total RNA from E11.5 FVB mouse embryos is extracted, mRNAs are purified, and the mRNA is fragmented to 100-200 nt long fragments. Refolded RNA fragments are used as input for IP and Flag elution. After yeast rRNA depletion from eluted RNAs, Illumina sequencing libraries of the ribosome-bound mRNA fragments are generated to identify hES9S-specific mRNA elements. IVT, *in vitro* transcript.

**Figure 3.**
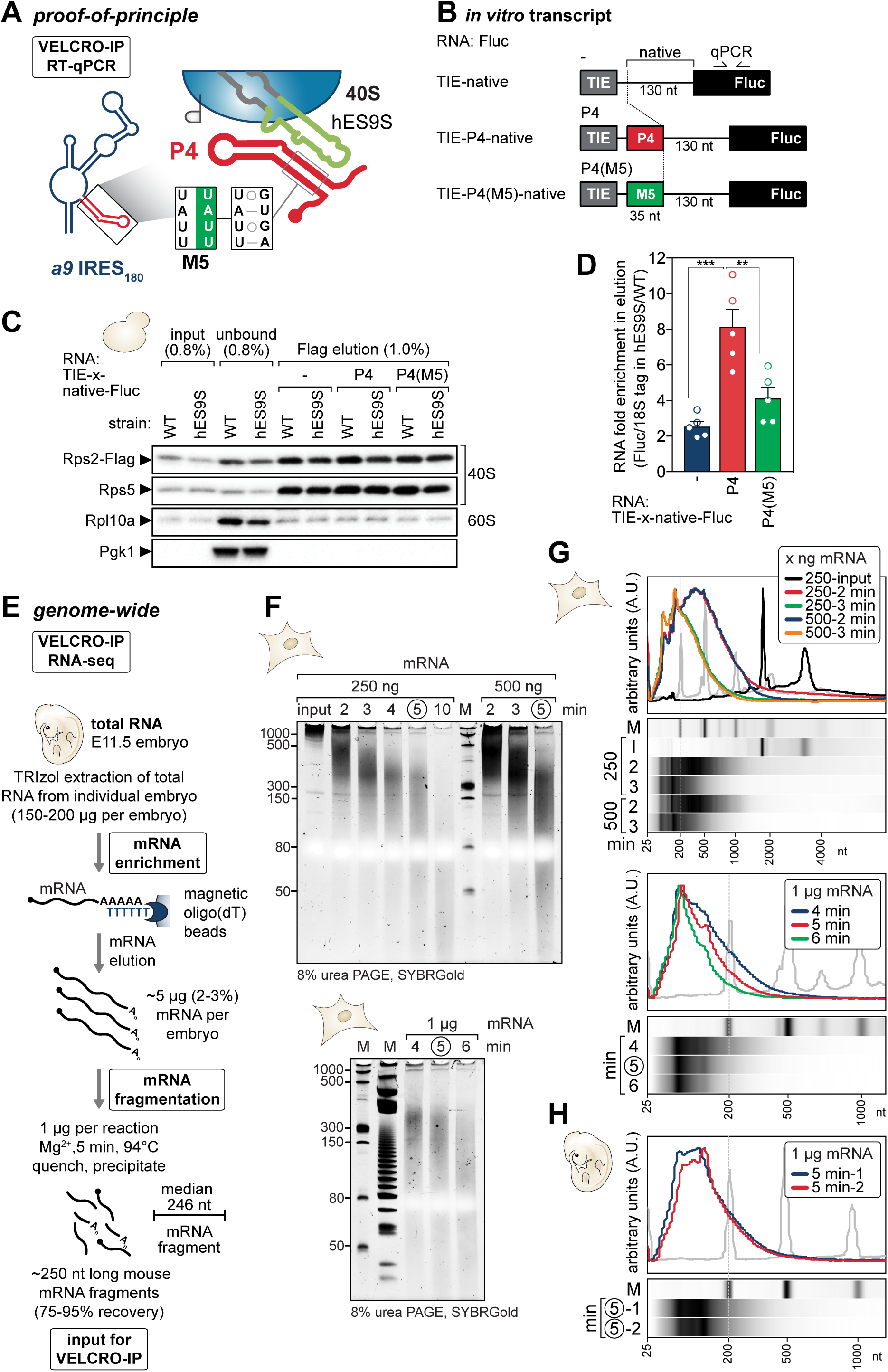
VELCRO-IP RT-qPCR serves as a proof-of-principle and mouse embryo mRNA fragmentation. (A) VELCRO-IP RT-qPCR: A zoomed-in view on the potential interactions between mRNAs and ESs, here hES9S binding to the *Hoxa9* P4 stem-loop (Leppek et al., 2020) or other target 5’ UTRs, that can be identified by the VELCRO-IP approach. The 4-nt inactive P4 mutant M5 (P4(M5)) serves as a negative control. (B) *In vitro* transcripts are generated using reporter mRNA plasmids as templates (see (Leppek et al., 2020)). The IVT RNAs contain the native spacer (-, negative control), P4-native (P4) or P4(M5)-native (P4(M5), P4-specific negative control) embedded in flanking constant regions (5’ TIE and 3’ Fluc ORF sequence). IVT RNAs have a total length of 475-510 nt and the Fluc ORF portion can be used for qPCR amplification across the three IVT RNA constructs. (C) WB analysis of same volumes of lysate (input), unbound fraction, and Flag peptide-eluted protein from beads to monitor ribosome enrichment of tagged (Rps2-Flag) and untagged (Rps5) 40S and 60S (Rpl10a) components in IVT RNA samples in combination with WT and hES9S yeast ribosomes. Cytoplasmic enzyme Pgk1 served as a negative control. The fraction loaded of input, unbound, and elution samples is expressed as percentage of the original lysate volume. Representative of n = 5 is shown. (D) Analysis of total RNA in the Flag peptide-elution by RT-qPCR using the same volumes of RNA per sample for the RT. Fluc transcript enrichment was assessed by internally normalizing Ct values to that of the respective 18S rRNA tag which controls for ribosome-IP efficiency per sample. We then compared respective hES9S to WT samples to assess specific RNA fold-enrichment of IVT RNAs. Average RNA fold enrichment ± SEM, n = 5. See also **Figure S3A-C**. (E) Schematic representation of embryo mRNA fragmentation for VELCRO-IP RNA-seq. In brief, total RNA extraction of stage E11.5 mouse embryos yields 2-3% of mRNA isolated on oligo(dT) beads, which is fragmented with magnesium ions to a length of 100-200 nt, and overall recovers <75% of input mRNAs as fragments. (F) Fragmented mouse mRNA from C3H/10T1/2 cells from different amounts (250 ng, 500 ng, and 1 µg) and timepoints of fragmentation alongside the 250 ng mRNA input were analyzed by 8% denaturing urea PAGE and visualized by SYBR Gold. The Low Range ssRNA Ladder (NEB) and the 20 bp Bayou DNA Ladder (Bayou Biolabs) were loaded for reference. (G) Fragmented mouse mRNA from C3H/10T1/2 cells from different amounts (250 and 500 ng) and timepoints of fragmentation (2 and 3 min) and the 250 ng mRNA input were analyzed on a mRNA Pico Chip (Agilent) on a Bioanalyzer (Agilent) and a zoomed-in view of the Bioanalyzer quantification and electronic gel analysis is shown. The marker (M, grey) is overlaid for reference. See also **Figure S3D**. (H) Fragmented mouse mRNA from stage E11.5 FVB embryos of 1 µg aliquots fragmented for 5 min at 94°C from two independent repeats of embryo harvest, RNA isolation, mRNA purification, and fragmentation (1, 2) is presented which was considered optimal to yield fragments of 100-200 nt in size. RNAs were analyzed on a mRNA Pico Chip (Agilent) on a Bioanalyzer (Agilent) and a zoomed-in view of the Bioanalyzer quantification and electronic gel analysis is shown. The marker (M, grey) is overlaid for reference. See also **Figure S3D**.

**Figure 4.**
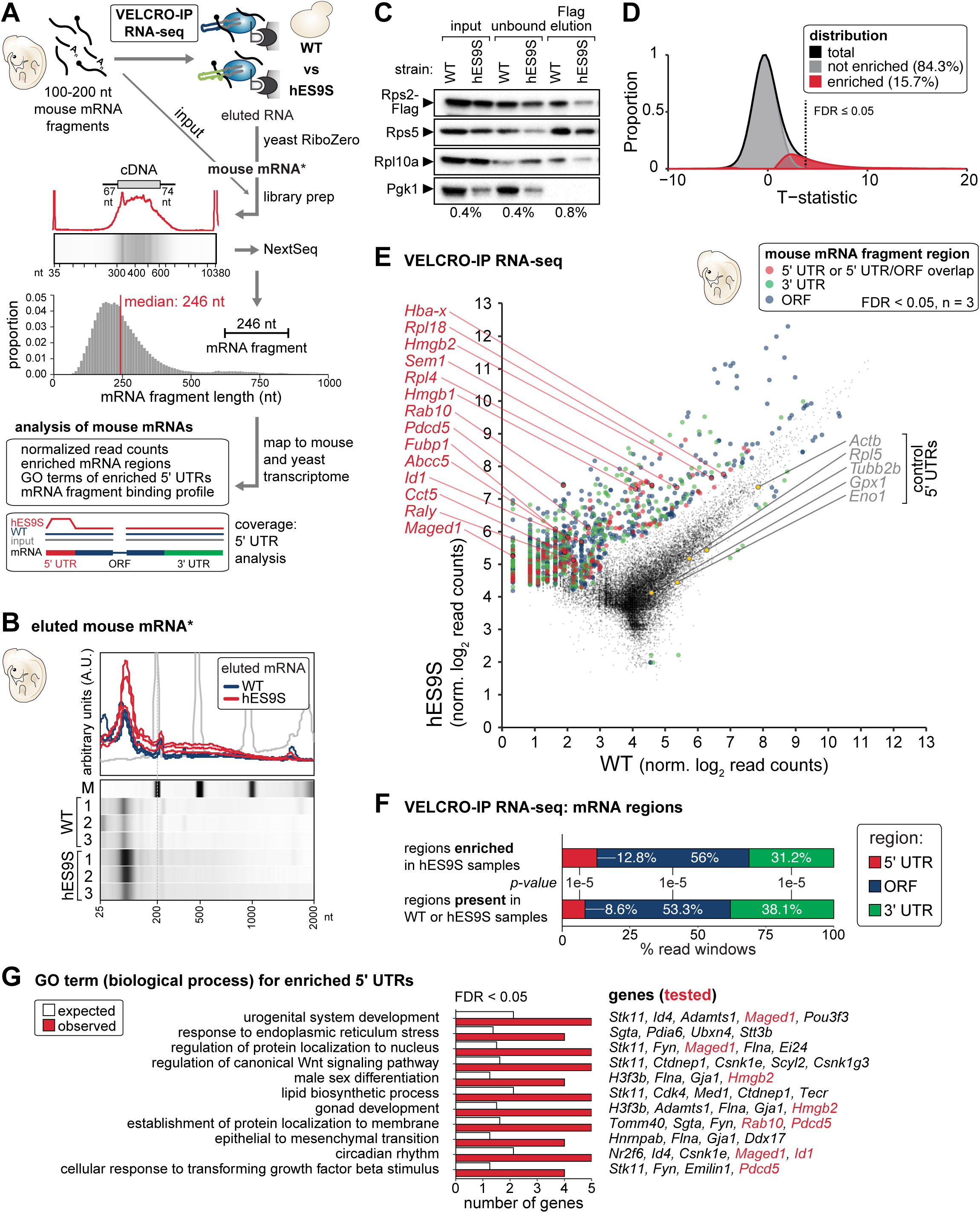
VELCRO-IP RNA-seq identifies global ES-mRNA interactions with positional resolution on mRNAs. (A) For VELCRO-IP RNA-seq, mRNA was isolated from stage E11.5 mouse embryos, fragmented to 100-200 nt and used as input for the IP. Total RNA obtained from all samples after elution and yeast rRNA depletion obtains ribosome-bound mouse mRNA that was subjected to library preparation and Illumina high-throughput sequencing (NextSeq). We included an mRNA fragment input sample for reference. The distribution of mRNA fragment lengths for all sequenced libraries is plotted and the median fragment length is 246 nt. We then map all reads to the mouse and yeast transcriptomes and only further analyze reads exclusively mapping to mouse mRNAs. (B) Eluted and yeast rRNA-depleted mouse RNA from three independent replicates of WT and hES9S VELCRO-IP experiments were analyzed on a mRNA Pico Chip (Agilent) on a Bioanalyzer (Agilent) and a zoomed-in view of the Bioanalyzer quantification and electronic gel analysis is shown. The marker (M, grey) is overlaid for reference. See also **Figure S3E**. (C) WB analysis as in Figure 3C to monitor efficient IP of 40S ribosomes for RNA-seq of mouse mRNA fragments. Representative of n = 3 is shown. (D) Kernel density of the distribution of t-statistics for test of differential enrichment of mRNA fragments bound to hES9S vs WT ribosomes is plotted in black. Empirical estimate of the decomposition of the distribution to null and non-null tests are plotted in grey and red, respectively. Dotted line indicates where the local FDR of 0.05 is. (E) RNA-seq results of independent replicates (n = 3) for each WT and hES9S samples. Normalized log read counts are presented for WT and hES9S-enriched mRNA fragments. Fragments less than FDR < 0.05 are colored according to the region in the mRNA they map to (see legend). We highlight fragments mapping to 5’ UTR or overlapping 5’ UTR/ORF (red) as well as 3’ UTR (green) and ORF (blue). We label mouse genes for which we identified enriched fragments in the 5’ UTR and/or 5’ region of the ORF and for whose 5’ UTRs we performed validation experiments. We further mark five control 5’ UTRs that are equally bound to both WT and hES9S 40S subunits and served as negative controls. See also **Figure S4, 5; Table S4**. (F) Analysis of regions mapping to the 5’ UTR, ORF or 3’ UTR in hES9S-enriched samples compared to their presence in WT or hES9S samples, each n = 3, expressed as % of total read windows identified. p denotes Indicated p-value is calculated by chi-square test. (G) Gene ontology (GO) analysis for biological process of total 87 5’ UTR regions (FDR < 0.05, n = 3) enriched with hES9S. Displayed are the expected and observed frequency of genes in categories meeting the cutoffs described in the methods. mRNA fragments present in either WT or hES9S samples were used as the background population (expected frequency). The presented genes in the eleven GO terms correspond to only 5’ UTR regions of hES9S-enriched mRNAs (FDR < 0.05, n = 3). Also see **Figure S6** for GO terms of ORF, 3’ UTR, and of the full mRNA (all regions) and **Table S5**.

**Figure 5.**
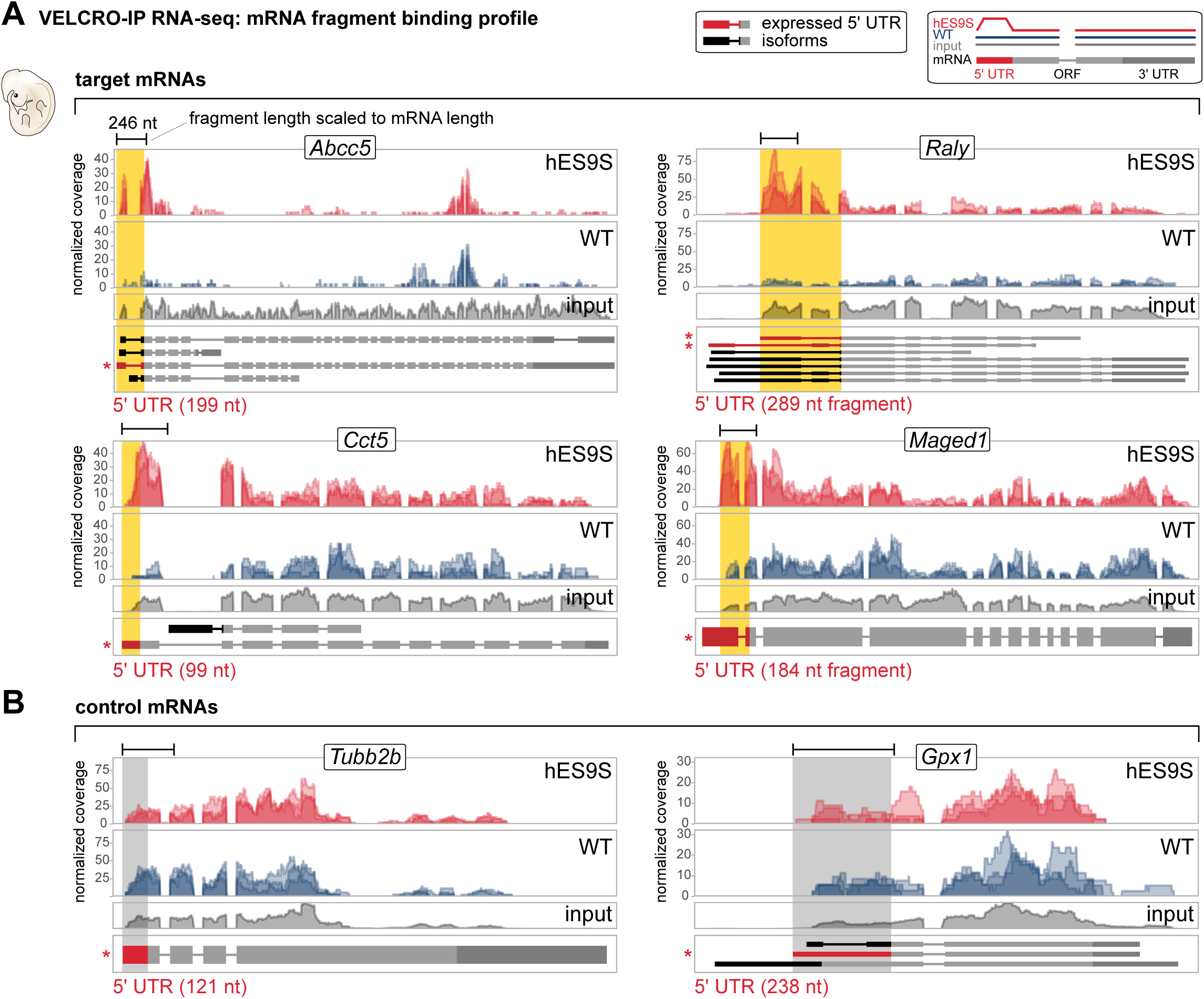
VELCRO-IP RNA-seq identifies hES9S-interacting 5’ UTRs with positional precision. (A) mRNA binding profile as coverage plots for four genes whose 5’ UTR-overlapping windows are significantly enriched in the hES9S over WT samples (FDR < 0.05, n = 3). Normalized per base coverage of individual biological replicate libraries for WT (blue) and hES9S (red) samples is plotted (above). All mRNA isoforms annotated in ENSEMBL are displayed below. Exon lengths are to scale while intron lengths are pseudo-scaled. The read coverage of the input mRNA fragments (grey) are also plotted for reference. 5’ UTR regions for the most likely expressed mRNA isoform in embryos is highlighted in red and the corresponding regions in the tracks is shaded in yellow. The 5’ UTR region picked for further experimental validation corresponds to the asterisk-marked isoform. The mRNA fragment length for each gene is scaled according to the mRNA length for the individual genes presented. See also **Figure S7A**. (B) The same analysis as in (A) was performed for two control 5’ UTRs that we found to equally bind to either WT or humanized 40S. 5’ UTR regions for the most likely expressed mRNA isoform in embryos is highlighted in red and the corresponding regions in the tracks, that also indicate no specific enrichment, is shaded in gray. See also **Figure S7B**.

**Figure 6.**
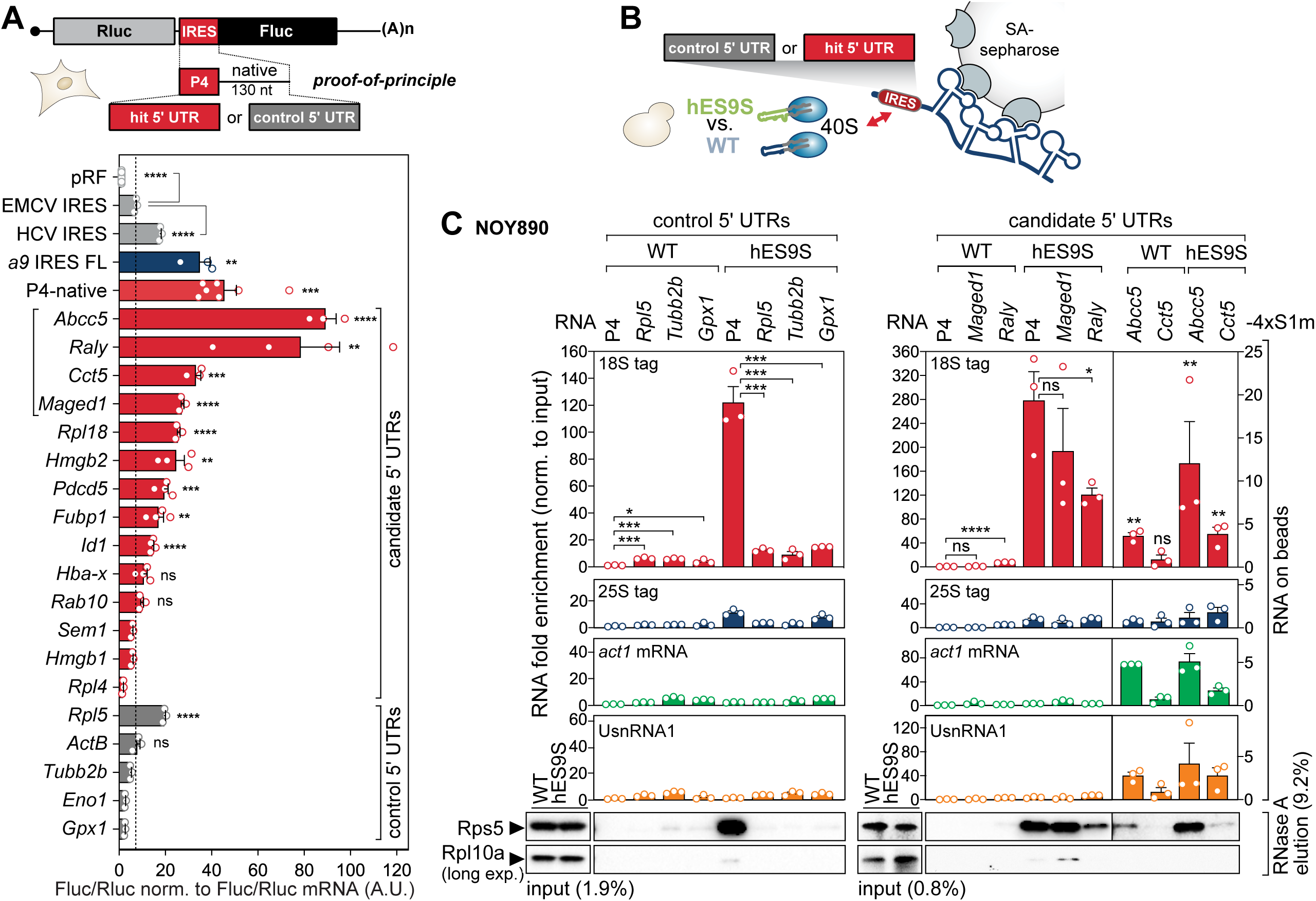
VELCRO-IP RNA-seq identifies hES9S-interacting 5’ UTRs with cap-independent initiation activity. (A) Based on the analysis in Figure 4, full 5’ UTRs as annotated in the ENSEMBL database were extracted for experimental validation. Bicistronic mRNA reporter genes containing no insert in the intergenic region (pRF, vector), candidate or control 5’ UTR sequences in the intergenic region were transiently transfected into mouse C3H/10T1/2 cells. Cells were harvested after 24 hours and cells from the same transfection were split in half for protein lysates and total RNA extraction, and subjected to luciferase activity measurement and RT-qPCR analysis, respectively. Relative luciferase activity is expressed as a Fluc(IRES)/Rluc(cap-initiation) ratio normalized to respective Fluc/Rluc mRNA levels, for the integrity of the bicistronic reporter mRNA to support cap-independent initiation activity of candidate 5’ UTRs, and expressed as average cap-independent initiation activity ± standard error of the mean (SEM), n = 3-8. pRF serves as negative control, the EMCV and HCV IRESs as IRES controls, and the full-length *Hoxa9* IRES-like element and P4-native as *Hoxa9* IRES-like references, respectively. EMCV IRES activity was used as a cutoff to assess candidate 5’ UTR cap-independent initiation activity. *a9* IRES FL: FL, full-length. (B) Schematic of the 4xS1m pulldown to probe the interactions of control and candidate 5’ UTR-4xS1m *in vitro* transcribed RNAs with WT and hES9S yeast ribosomes. Pre-coupled 5’ UTR-4xS1m RNA on SA-sepharose beads are incubated with lysates of WT and hES9S yeast strains (NOY890). Ribosome-RNA RNP enrichment *in vitro* is monitored by RT-qPCR for tagged rRNA and other RNA classes normalized to the input, and WB analysis for RPs. (C) 4xS1m pulldown to probe the interactions of candidate 5’ UTR-4xS1m *in vitro* transcribed RNA with WT and hES9S yeast ribosomes. Three control 5’ UTRs served as negative controls and the four candidate 5’ UTRs that display highest cap-independent initiation activity in (A) were tested using *Hoxa9* P4 as a reference. Pre-coupled 5’ UTR-4xS1m RNA on SA-sepharose beads are incubated with yeast lysates to form ribonucleoproteins (RNPs) *in vitro* and protein and RNA are specifically eluted. After the formation of ribosome-RNA RNPs *in vitro*, beads are split in half: total RNA is eluted with TRIzol, and protein is eluted with RNase A. rRNA bound to the 4xS1m- fused RNA is quantified with primers specific for 18S and 25S rRNA tags (RNA on beads). The P4 serve as a positive control. To indicate specific enrichment of RNA, fold enrichment of RNAs was determined by RT-qPCR using same volumes of eluted RNA and normalizing Ct values of each sample to their respective RNA input (WT or hES9S). Yeast *actin* (*act1*) and yeast UsnRNA1 serve as negative controls for an mRNA and a non-coding RNA, respectively. Ribosome enrichment, particularly ribosomal proteins of the 40S and 60S subunit, was assessed by WB analysis of same volumes of protein released from beads by RNase A. The fraction loaded of input and elution samples is expressed as percentage of the original lysate volume. The P4-4xS1m/WT sample was used to normalize for RNA fold enrichment (set to 1). Average RNA fold enrichment, SEM, n = 3; ns, not significant; long exp., long exposure. See also **Figure S7C**.

## RESULTS

### Engineering of yeast ribosomes with customized rRNA expansion segments for VELCRO-IP

Based on our previous finding that ES9S directly recruits 40S ribosomal subunits to the 5’ UTR of a key developmental gene, *Hoxa9* (Leppek et al., 2020), we utilize humanized ES9S as the proof-of-principle example in the development of our method to identify the mammalian mRNA interactome of ESs (**Figure 1, S1**). Mouse ES9S and human ES9S are 100% identical, and while we refer to chimeric ribosomes as humanized ES9S (hES9S), the *Hoxa9* 5’ UTR as well as the transcriptome employed throughout this study is of mouse origin.

When the secondary structures of 18S rRNAs for the evolutionary distant baker’s yeast *Saccharomyces cerevisiae* (*S. cerevisiae*) (Armache et al., 2010) and human (*H. sapiens*) (Natchiar et al., 2017) are compared (**Figure 1A; S1A, B**), the basal stem region of helix h39 adjacent to ES9S is highly conserved while the distal portion of ES9S is highly variable in length, structure and sequence (**Figure 1A-C**, boxed region). Even among vertebrate species that are more closely related, such as chicken (*Gallus gallus*), axolotl (*Ambystoma mexicanum*), frog, (*Xenopus laevis*) and zebrafish (*Danio rerio*), there are nucleotide insertions and deletions in ES9S that affect ES9S structure (**Figure 1C; S1D, E**). Their presence was confirmed by RT-PCR using cDNA from tissues of the respective species (**Figure 1B, C**).

This divergence in ES sequence is the prerequisite for and essence of VELCRO-IP. We harness the interspecies variability in ESs to uncover the differential mRNA interactome of a defined ES and to functionally test the importance of ES sequences for species-specific mRNA-binding and translation (**Figure 1D, E**). VELCRO-IP is designed to be applicable to any ES using the ribosome engineering system in yeast (**Figure 1E**).

The best strategy of isolating functional mRNA-ES binding events is in context of the mature ribosome, one of the most complex cellular RNPs. Thus, we designed VELCRO-IP to rely on the constant core of the ribosome, with all its exposed binding surfaces for proteins and RNAs, editing only one ES sequence at a time. This relies on several crucial strategies in terms of ribosome design: 1. Employing yeast ribosomes as “minimal” ribosomes onto which evolutionary distant metazoan ES sequences can be scarlessly transplanted; 2. Carefully designing interspecies ES transplants according to rRNA structure such that highly conserved constant regions are chosen as the edit site; 3. Inserting RNA sequence tags into 18S and 25S rRNA to distinguish edited rRNA-ribosomes from WT ribosomes for IP-enrichment analysis by RT-qPCR; 4. Generating tagged WT ribosome strains containing the yeast ES along with chimeric ribosome strains for direct comparison; 5. Tagging the 40S ribosomal subunit with a Flag tag by using a rDNA deletion yeast strains that contain endogenously C-terminally Flag-tagged RPS2/uS5 to isolate WT and chimeric 40S ribosomal subunits. This approach yields yeast ribosomes that contain 18S and 25S rRNA sequence tags, a Rps2-Flag tag, and either a WT or chimeric ES of choice (**Figure 1E**).

We first engineered chimeric ribosomes in yeast by “humanizing” yeast 18S rRNA exclusively in the distal part of ES9S, which is divergent between the two species. VELCRO-IP uses the yeast ribosome core to accomplish this, because yeast only has a single rDNA locus containing hundreds of tandemly repeated rDNA copies. The entire rDNA locus can be deleted and complemented with exogenous expression plasmids containing engineered rDNA sequences (Nemoto et al., 2010; Wai et al., 2000). Such engineered “humanized” hybrid ribosomes for ES9S (**Figure 1E, S1**) contain hES9S introduced scarlessly into the h39 stem region of yeast 18S rRNA that is highly conserved in sequence and structure (boxed region in **Figure 1A**). We found it crucial to design hybrid rRNAs according to RNA structure, only transplanting the most distal part of the foreign ES onto yeast 18S rRNA (**Figure S1B, C**). This complementary exchange of smaller regions is important since deletion of large regions from most ESs can lead to ribosome biogenesis defects and to severe viability defects (Jeeninga et al., 1997; Ramesh and Woolford, 2016; Sweeney et al., 1994). Furthermore, we introduced unique sequence tags into both 18S and 25S rRNAs (Fujii et al., 2009) to quantitatively distinguish the humanized chimeric ribosomes from potentially remaining untagged wildtype (WT) ribosomes by RT-qPCR prior to ribosome purification (**Figure 1E, S2**). As VELCRO-IP relies on the comparison of the interactomes of chimeric versus WT ribosomes, we in parallel generated tagged but otherwise unmodified ribosomes that retain yeast ES9S (referred to hereafter as WT). For tagged hES9S-ribosome containing yeast strains, we confirmed that yeast cells that are induced to exclusively contain tagged ES9S-humanized ribosomes are viable, and only show a slight growth defect in comparison to tagged WT rRNA-containing cells in a viability assay (Leppek et al., 2020).

This paved the way for the successful isolation of yeast strains after rDNA plasmid shuffling into the NOY890/Rps2-Flag strain that solely contain plasmid-derived tagged hES9S or WT 18S rRNA-ribosome (**Figure 1E, S2A**) (Nemoto et al., 2010). Positive clones after shuffling were characterized by RT-PCR specific for the length difference of the ES9S sequence and the presence of the 18S rRNA tag (**Figure S2B**), as well as the number of tagged to endogenous ribosomes present after shuffling in these cells (**Figure S2C**). To this end, we endogenously C-terminally Flag-tagged *RPS2/uS5*, which has previously been used to successfully tag yeast ribosomes for isolation (Jan et al., 2014) (**Figure S2D**). Sucrose gradient fractionation of yeast lysates and Western blot analysis confirmed that Rps2-Flag protein is present in the heavy polysomes in both the Flag-tagged WT and hES9S-humanized ribosome strains (**Figure S2D**). This lack of difference in polysome profiles indicate no difference in translation rates between the strains. Additionally, the comparison of hES9S and WT strains with and without the Rps2-Flag indicated that another control 40S ribosomal subunit component, Rps5/uS7, is found normally incorporated into translating polysomes in both strains. Rps2-Flag is present in the same heavy polysome fractions as Rps5. Flag-tagged RP incorporation into translating ribosomes is a prerequisite for isolation of mature ribosome isolation by VELCRO-IP. These strains could therefore next be used as a tool to study species-specific mRNA-ES interactions.

### VELCRO-IP employs purification of engineered humanized yeast ribosomes

With straightforward generation of tagged chimeric ribosomes on hand, we next asked whether and which mammalian mRNAs in the transcriptome may recruit the 40S ribosome by binding to hES9S. To answer this question, we designed a technology that combines a pulldown strategy to first capture chimeric and WT yeast ribosomes and then to use the captured ribosomes as bait to identify differentially bound mRNAs genome-wide (**Figure 2**). We termed this strategy VELCRO-IP (<u>v</u>ariable <u>e</u>xpansion <u>s</u>egment-ligand <u>c</u>himeric <u>r</u>ib<u>o</u>some-immuno<u>p</u>recipitation). The modularity of this workflow allows the choice of not only the ES but also the choice of any tissue- or cell-derived transcriptome that is relevant to a biological question.

We first sought to confirm the successful isolation of a known, physiologically relevant interaction between an *in vitro* transcript of a *Hoxa9* 5’ UTR element and the hES9S chimeric ribosomes. For this, we devised a proof-of principle approach, VELCRO-IP RT-qPCR (**Figure 2**, left), that lays the groundwork for transcriptome-wide VELCRO-IP RNA-seq (**Figure 2**, right). The VELCRO-IP protocol for both scenarios is based on a series of carefully optimized biochemical steps.

First, the Flag-pulldown of ribosome-mRNA complexes from yeast lysates was performed to enrich for 40S ribosomal subunits. For this, WT and hES9S-rRNA expressing NOY890/Rps2- Flag strains were harvested in mid-log phase of actively translating cells. To ensure scalability, lysates were prepared by powderizing cell pellets in liquid nitrogen using a mortar and pestle. As mRNA input, either *in vitro* transcribed RNA from plasmids encoding the *Hoxa9* 5’ UTR (proof-of-principle) or fragmented poly(A)-enriched mRNA from stage E11.5 mouse embryos (genome-wide) was used, as described in further detail below. For ribosome isolation, RPS2-Flag tagged 40S ribosomes were immuno-precipitated from lysates on anti-Flag M2 affinity agarose gel. Previous experience had shown that agarose gel is advantageous over magnetic beads to cleanly isolate ribosomes (Simsek et al., 2017). This first purification step yields a ribosome-resin of washed 40S ribosomal subunits bound via Rps2-Flag before incubation with an RNA input source. For VELCRO-IP RNA-seq, fragmented mRNA from E11.5 mouse embryos were pooled and refolded in three steps of decreasing temperature to slowly reconstitute RNA structures such as short stem-loops. An input RNA sample was also collected at this step for RNA sequencing. Refolded RNA was added to WT and hES9S-ribosome-coupled beads for IP and bound ribosome-mRNA fragment complexes were washed and eluted off the anti-Flag beads using competitive 3xFlag peptide elution. The IP- and elution-efficiency was monitored by protein analysis by WB and total RNA extraction for RNA seq library preparation. A key design element that ensures the specificity of the detected hES9S-mRNA interaction in this protocol is the parallel generation and comparison of the interactomes of the WT and hES9S yeast ribosomes. This workflow can be performed in a day and is highly modular as it relies on sequential steps of 1. bead-based ribosome purification, 2. incubation with any pool of putatively interacting RNAs, 3. efficient, ribosome-specific Flag peptide elution, and 4. quantitative analysis of the eluted RNA.

### VELCRO-IP RT-qPCR enables interrogation of variant ES-specific ribosome-mRNA interactions

We have previously shown that hES9S ribosomes are sufficient to reconstitute binding to the *Hoxa9* 5’ UTR, particularly to the 35 nt stem-loop P4 in the *Hoxa9* IRES-like RNA element, which highlights the ES specificity of this mRNA-rRNA binding event (**Figure 3A**, (Leppek et al., 2020)). P4 thus served as an ideal positive control. In a proof-of-principle experiment, we optimized 40S ribosome isolation from yeast lysates and used *in vitro* transcripts containing P4. This P4 positive control, labeled P4-native, contains a native spacer sequence between the P4 and the start codon that is required for its function. The native spacer sequence without P4 thus serves as a negative control. In addition, our previous data also provide another negative control in the form of a 4-nt inactivating mutation within P4, termed P4(M5), that disrupts the ES interaction. Both the native spacer alone as well as P4(M5)-native are our negative controls. All three RNA constructs were used to test the specificity of hES9S-ribosome interactions with P4 (**Figure 3B, S3A, B**). For the pulldown, ∼500 nt long *in vitro* transcripts of native, P4-native or M5-native RNAs were generated that were flanked by constant sequences, stemming from mRNA reporter previously used (Leppek et al., 2020), that allow for comparable Fluc-specific qPCR quantification (**Figure S3B**). Purified yeast 40S ribosomes on Flag beads were incubated with these RNA transcripts, and washed RNA-ribosome complexes were eluted with 3xFlag peptide. The analysis of specific protein and RNA enrichment in eluates demonstrated that (1) VELCRO-IP cleanly isolates tagged 40S ribosomal subunits (**Figure 3C**) and that (2) in comparison to WT, yeast hES9S-ribosomes enrich P4-native-containing transcripts about 4-fold more than native-spacer RNA (**Figure 3D, S3C**). The clear reduction in hES9S-ribosome binding to the inactive P4(M5) mutant highlights the specificity and sensitivity of the VELCRO-IP approach.

### VELCRO-IP RNA-seq uses mRNA fragments to map hES9S-interacting mRNA regions

The VELCRO-IP RT-qPCR results for control *in vitro* transcripts paved the way for a genome-wide version of the ribosome pulldown experiment, VELCRO-IP RNA-seq, that uses fragmented mouse embryo mRNAs to identify mRNA regions that preferentially rely on hES9S for ribosome binding (**Figure 2**, right). To gain such positional information of bound mRNA regions an ES preferentially interacts with – a localized window, for example, within 5’ UTR, ORF or 3’ UTR (**Figure 1D**) – we decided to use random fragments of the input mRNA in the size range of 100-200 nt (**Figure 3E**), a size allowing P4-like stem-loops of at least 35 nt to fold (**Figure 3A**). To generate a pool of endogenous mouse embryo mRNAs as a physiological source of RNA, up to 10 stage E11.5 mouse embryos were harvested individually which yielded a 150-200 µg total RNA per embryo. From total RNA, poly(A) mRNA was isolated on magnetic oligo(dT) beads, which yielded ∼5 µg mRNA (2-3% of total RNA input) per embryo. Purified embryo mRNA was fragmented to 100-200 nt size range by hydrolysis with magnesium ions and heat. Fragmentation was optimized for time (0-10 min) and mRNA input amount (250 ng, 500 ng and 1 µg mRNA) monitoring RNA size by urea-PAGE (**Figure 3F**) and by Bioanalyzer analysis (**Figure 3G, H; S3D, E**). mRNA fragmentation was optimized using mRNA from C3H10T1/2 mesenchymal cells which performed identically when we employed purified stage E11.5 embryo mRNA (**Figure 3G, H; S3D, E**). Immediate precipitation recovered 75-95% of input mRNA as mRNA fragments. VELCRO-IP RNA-seq uses 10 µg of fragmented mRNAs as input. After yeast ribosome-IP, ribosomes were incubated with fragmented and refolded mouse embryo mRNA and ribosome-mRNA complexes were eluted with Flag peptide (**Figure 4A**). Eluted RNAs are mainly comprised of yeast rRNA which were depleted to increase the representation of mouse mRNA fragments in the final RNA-seq library. We detected an overall enrichment of mouse mRNA fragments in hES9S samples compared to WT controls by Bioanalyzer analysis (**Figure 4B**). The IP and elution efficiency were confirmed by WB analysis (**Figure 4C**). The sequencing libraries were prepared from yeast rRNA-depleted eluted RNAs, using randomly primed reverse transcription and incorporating unique molecular tags prior to amplification. The cDNA libraries were sequenced using the high-throughput Illumina platform (**Figure 4A**).

### VELCRO-IP RNA-seq identifies ES9S-interacting mRNA elements genome-wide

We performed and sequenced three replicate VELCRO-IP RNA-seq experiments each for WT and hES9S. The three independent repeats for WT and hES9S samples were highly reproducible (**Figure S4**). The final median fragment length observed in the sequencing library was 246 nt (**Figure 4A**). We counted sequencing reads in 200 nt windows across the genome and detected 18989 windows across 2610 genes with sufficient coverage for statistical tests of differential enrichment of mRNA fragments bound to hES9S- over WT yeast ribosomes. Using an empirical modeling of the test statistic distribution, we estimated that 15.7% of the 18989 regions are differentially enriched and thus have binding dependency on hES9S (**Figure 4D**). At the false discovery rate (FDR) of 5%, we are able to confidently classify 1491 regions over 460 different genes as strong candidates for further analysis (**Figure 4E; S5; Table S4**). This indicates a pervasive, hES9-dependent binding of mRNA regions to ribosomes throughout the transcriptome.

We did not observe an overwhelming overrepresentation of the hES9-enriched mRNA regions in a particular 5’ UTR/ ORF/ 3’ UTR annotation category, though we detected a slight overrepresentation of reads mapping to the 5’ UTR over ORF/ 3’ UTR (∼1.7-fold, p<1×10^-5^; **Figure 4E, F**). Among the group of 460 genes whose mRNAs bound to humanized ribosomes (**Table S4**), 87 genes were identified whose enriched regions overlap with the 5’ UTR. They are enriched for Gene Ontology (GO) terms involving developmental and differentiation processes – for example, regulation of Wnt signaling pathways, gonad development, and urogenital system development (**Figure 4G, Table S5**). Another interesting category is that of circadian rhythm, whose biology frequently involves translational control for temporal expression patterns, for example gene <u>m</u>elanoma <u>a</u>ntigen-encoding <u>ge</u>ne D1 (*Maged1*) and inhibitor of DNA binding 1 (*Id1*)), which we later included in validation experiments. GO term enrichment analysis for CDS, 3’ UTR, or all regions together further revealed other diverse types of functional annotations such as cell cycle, DNA damage responses or muscle contraction (**Figure S6; Table S5**). These data together suggest that hES9S-bound mRNAs may be involved in post-transcriptional regulation of multiple important functional pathways, especially in mammalian embryonic development.

### hES9S-interacting mRNA elements mediate cap-independent translation

In order to functionally validate identified hES9S-mRNA interactions, we focused on the most significantly enriched mRNA 5’ UTR regions (**Figure S5**). We based this direction on a confirmed function we had previously discovered: our prime example of the P4 stem-loop in *Hoxa9* 5’ UTR that interacts with hES9S to recruit the 40S ribosomal subunit for translation initiation (Leppek et al., 2020). Thus, we next asked if hES9S-enriched 5’ UTRs mediate cap-independent activity, similar to that of the *Hoxa9* 5’ UTR. We chose the 14 most significantly hES9S-enriched 5’ UTRs as candidates (red in **Figure 4E**, **S5**), and tested their full-length mouse 5’ UTRs in bicistronic mRNA reporters. The selection of the 5’ UTR isoform sequence was guided by the expression profile of the candidate mRNAs in the input and their mRNA binding profile of enriched fragments in hES9S samples (**Figure 5A, S6A**). For some shorter (<246nt, median fragment length) candidate 5’ UTRs in which the enrichment peak extends into the ORF, we restricted our analysis to the 5’ UTR. 9 out of 14 candidate 5’ UTRs exhibit cap-independent activities higher than the viral encephalomyocarditis virus (EMCV) IRES, which served as a reference and positive control (**Figure 6A**). Importantly, to assess the likelihood of cap-independent activity in 5’ UTRs without hES9S interaction, we also tested five control 5’ UTRs that clearly were not enriched in hES9S over WT in our genome-wide VELCRO-IP RNA-seq data. They were selected based on their estimated negative predictive values and confidence intervals (**Figure 4E, 5B; S6B**). These results suggested that the specificity of the mRNA-hES9S interaction as determined by our VELCRO-IP RNA-seq functionally selected for cap-independent activity (**Figure 6A**).

Beyond the functional correlation with cap-independent translation initiation, an orthogonal approach was employed to further validate the interaction of the 5’ UTRs with chimeric hES9S-ribosomes for four of the tested candidates with the strongest cap-independent initiation activities: ATP binding cassette subfamily C member 5 (*Abcc5*), hn<u>R</u>NP protein <u>a</u>ssociated with lethal <u>y</u>ellow (*Raly*), chaperonin containing TCP1 subunit 5 (*Cct5*), and *Maged1.* Three negative control 5’ UTRs were also included: *Rpl5*, tubulin beta 2B class IIb (*Tubb2b*), and glutathione peroxidase 1 (*Gpx1*). In a reverse approach to VELCRO-IP, we performed 4xS1m pulldown experiments that we had established previously (Leppek and Stoecklin, 2014; Leppek et al., 2013), using the 5’ UTRs as RNA bait for WT and hES9S-ribosomes using yeast cell lysates as input (**Figure 6B**). Compared to our positive control, *Hoxa9* P4, there is no enrichment for any of the control 5’ UTR, including *Rpl5*, to hES9S-ribosomes. In a stark contrast, we observe significant enrichments of all candidate 5’ UTRs identified in our VELCRO-IP RNA-seq experiments (**Figure 6C, S6C**). Noticeably, *Maged1* and *Raly* bind to hES9S 40S ribosomal subunit in the same range as the *Hoxa9* P4 element. These results additionally demonstrate the high specificity of our genome-wide VELCRO-IP RNA-seq analysis.

*Maged1* is known to be important for brain and bone formation (Bertrand et al., 2004; Liu et al., 2015), including possible regulation of homeodomain transcription factors such as *Dlx5* and *Msx2* (Masuda et al., 2001). *Raly* encodes a RNA binding protein, which has been implicated in early pre-implantation embryonic development (Michaud et al., 1993). These data thus identified critical physiological regulators that specifically recruit ribosomes for cap-independent translation through hES9S. Interestingly, prior comparative analysis of *Maged1* expression during brain and embryonic development has revealed a discrepancy between mRNA and protein expression levels, suggesting that its expression levels are indeed regulated at the post-transcriptional level (Bertrand et al., 2004). Taken together, our VELCRO-IP RNA-seq approach represents a powerful tool to reveal how ribosome-mediated control of gene regulation is achieved at the molecular level in a genome-wide manner. In combination with orthogonal mRNA reporter and pulldown assays for validation, this methodology represents a targeted strategy to further identify mRNAs that directly bind to any other ES on the ribosome in a species-specific manner. This technology has revealed an additional, selective subset of mRNAs that require a specific ES sequence for binding to ribosomes. Further validation showed that these additional mRNAs undergo cap-independent translation initiation.

## DISCUSSION

### The unique advantage of VELCRO-IP

It has been historically challenging to assign functions for rRNA ESs. To the best of our knowledge, this persistent lack of knowledge about functions of ES even three decades after their discovery in the 1980s (Gerbi, 1986, 1996) is due to two reasons. On one hand is their missing structure: unlike the more rigid, universally conserved structure of the ribosome at its core (Melnikov et al., 2012; Taylor et al., 2009), ESs are notoriously challenging to visualize on the ribosome in cryo-EM structure reconstructions because they display structural flexibility and dynamics forming the outer shell of the ribosome (Armache et al., 2010). They are thus considered the “black box” of the ribosome in the field of structural biology. Beyond the missing structure, on the other hand, it further remains impossible to genetically manipulate ESs in most eukaryotic species. Eukaryotic rDNA consists of hundreds of tandemly repeated units per cluster, spread across multiple chromosomal loci. In humans, the ten rDNA clusters make up <0.5% of the diploid human genome but it remains mostly unclear how many of the clusters are transcribed at the same time (Denissov et al., 2011; Gerbi, 1986; Parks et al., 2018; Stults et al., 2009). However, rRNA transcription constitutes >80% of cellular RNAs with thousands of ribosomal subunits synthesized each minute in actively growing eukaryotic cells (Lewis and Tollervey, 2000; Warner, 1999), implying that at least a significant proportion of rDNA copies are likely to be functional and under selection for growth. Thus, to investigate an rRNA sequence feature like ESs that are specific to different species – for example, mammalian species like humans compared to yeast – the rDNA must be edited for hundreds of copies at once to gain a homogeneous ribosome population, against the selective pressure for fully functional rRNA copies. This remains a major barrier to functional investigation of ESs.

Until VELCRO-IP, which can directly address an ES function in isolation biochemically, previous methodologies for studying rRNAs have mostly remained observational. These include imaging-based approaches such as immune-gold staining (Miller and Beatty, 1969), metabolic labelling (Stefanovsky and Moss, 2016), or *in situ* hybridization with fluorescent probes (Cao et al., 2019) as well as ChIP-seq based methods (Mars et al., 2018; Zentner et al., 2011), which allow analysis of transcription and localization of rRNA clusters within species (McStay and Grummt, 2008). The rDNA operons may have variants that could correlate with a function – for example in prokaryotes, such variant rDNA operons can be selectively expressed under stress (Kurylo et al., 2018; Song et al., 2019). In addition, analysis of large-scale human genome and transcriptome datasets have suggested differential expression of variant rDNA species across individuals and tissues (Parks et al., 2018). However, these methods cannot directly address cross-species sequence variation and function of ES. Furthermore, while ESs have been previously manipulated in yeast, they all yielded general function of ESs in ribosome biogenesis (Jeeninga et al., 1997; Ramesh and Woolford, 2016; Sweeney et al., 1994) but these complete deletions of ESs preclude a more specific analysis of ES functions beyond their role in cell growth and viability, such as in translational control. VELCRO-IP is the first technique that can directly and cleanly isolate and test biochemically the decades-old hypothesis of what molecular functions a variable ES may have in translation regulation by using scarless switching of ES sequences, rather than deletion of larger rDNA portions prone to interfere with accurate ribosome assembly and cell viability.

ES regions provide a playground for evolutionary diversity among rRNA sequences that differ immensely between species and even between tissues in the same organism (Parks et al., 2018). VELCRO-IP RNA-seq can distinguish eukaryotic-specific ES when transplanted onto a minimal yeast ribosomal core complex. We mainly detect species-specific indels for ES9S across eukaryotes (**Figure 1C**) that may be interesting to correlate with their respective mRNA binding profile. Another physiological example in vertebrates is presented in the distinct maternal-type and somatic-type ribosomes in zebrafish wherein the majority of detected differences can be attributed to ES sequence variability based on nucleotide substitutions and indels (Locati et al., 2017). It will be interesting to investigate the effect of such smaller ES changes, in contrast to the evolutionary distant human and yeast ES sequences, on ribosome function, and how much of their diverse interactome can be captured by VELCRO-IP that is based on differential affinity of *trans*-acting factors.

VELCRO-IP RNA-seq reveals species-specific mRNA binding for translation regulation, but how do diverse ES sequences arise and are maintained in the genome that might mediate unknown ribosome-linked functions encoded in ESs? The mechanisms by which a tandem array of rDNA genes is generated has been hypothesized to arise from unequal crossing-over or excision of an rDNA copy coupled with rolling circle replication and reintegration (Ganley and Kobayashi, 2007; Hourcade et al., 1973). Thus, rDNA is in a constant flux because point mutations and insertions or deletions in individual rDNA copies may occur randomly throughout the gene. Consequently, new variant repeats can gradually spread through the rDNA loci and through individual species. The fact that a conserved common core structure exists in ribosomes in all kingdoms of life, including bacteria, archaea, plants, and animals, suggests that the architecture of the primitive ribosome may have been established before the separation into distinct phylogenetic branches, and that any changes to the ribosome core are likely to be harmful and selected against. Therefore, negative selection is thought to drive down rDNA copies with a deleterious mutation while positive selection may favor the spread of useful changes. Such useful changes might be presented in species-specific variations in ES sequence that await testing by VELCRO-IP RNA seq.

### The biological significance of rRNA-mRNA interactions identified by VELCRO-IP RNA-seq

To date, mRNA-rRNA interactions have been mainly implicated for translational control in viruses or bacteria. In eukaryotes, one example of mRNA-rRNA interaction is found between the purine-rich sequence in the histone H4 mRNA coding region and helix h16 of the 18S rRNA, whose base pairing tethers the 40S ribosome to the start codon (Martin et al., 2016). These mRNAs contact conserved rRNA segments rather than ESs. It is unknown whether direct mRNA-rRNA interaction could be a globally utilized mechanism for translational control in eukaryotes.

Using hES9S chimeric ribosomes allowed us to define a class of ES9S-responsive mRNA 5’ UTRs. Aimed at revealing mRNAs harboring cap-independent ability to recruit the ribosome, our study discovered and functionally confirmed 5’ UTRs previously unknown cap-independent translation activity, which bind to the 40S ribosomal subunit via ES9S. Unexpectedly, our data also reveal additional hES9S-interacting mRNA fragments mapping to the CDS and 3’ UTR, whose functional contribution remain to be tested. Such RNA elements in these regions may mediate functions via the ribosome in diverse steps of RNA metabolism such as mRNA localization or decay, other than 5’ UTR interactions for translation initiation. Prior to this study, it has been an interesting but unanswerable question whether the massively expanded eukaryotic ESs on the surface of the ribosome could pose a landing pad for mRNAs for their recruitment to the ribosome. Our results provide evidence for selective recruitment of mRNAs by a specific ES of the eukaryotic ribosome on a genome-wide scale.

### Potential applications of VELCRO-IP for any rRNA ES

In the future, we foresee numerous applications of VELCRO-IP in probing the effects of rRNA ESs on translation regulation. The presented ribosome engineering in ES regions is a modular and versatile tool that can be adaptable site-specifically to any ES (**Figure 1D)**. We encourage to apply this method to ESs on both 40S as well as 60S subunits, by Flag tagging Rps2 or a large ribosomal protein. VELCRO-IP RNA-seq approach can potentially be utilized widely to study the multitude of other ESs on the ribosome across many different species of interest to reveal other classes of direct mRNA recruitment to the ribosome. This will help us to understand how the sequence variation in ES regions influence the evolutionary diversity in the expression of the translatome. Beyond ES-mRNA interactions, we foresee that VELCRO-IP can be adapted to investigate the ES-bound proteome by coupling it to mass spectrometry (MS), VELCRO-IP MS, to map the ES-specific binding sites of ribosome-associated proteins (RAPs) (Simsek et al., 2017) that comprise the outer protein-shell of the ribosome. For example, our lab has recently discovered a distinct function for the 25S rRNA ES27L in translation fidelity (Fujii et al., 2018) that originates from an ES-protein interaction on the 60S ribosomal subunit, whose co-complex with the ES has also been structurally investigated (Knorr et al., 2018; Wild et al., 2020). Moreover, there appear to be hundreds of additional RAPs that bind to the mammalian ribosome as part of the ‘ribointeractome’, which may employ specific ESs as binding sites (Simsek et al., 2017). It will be interesting to see if ES-mediated protein recruitment to the ribosome can endow the ribosome with novel functions unique to specific tissues or organisms.

Together, our species-specific engineering approach in yeast provides an elegant and robust solution to address ES-specific functions in context of the ribosome. VELCRO-IP in principle allows the study of any ES and its species-specific interactions with *cis*-regulatory mRNA elements or RAPs. We envision that this methodology will lead to a more precise understanding of rRNA function in gene regulation, in other translation-coupled cellular processes.

## LIMITATIONS

The design of a chimeric ribosome will likely require a minimum size of the replacement ES. The function of ESs in ribosomal biogenesis and export has been described mainly in 25S rRNA (Bradatsch et al., 2012; Greber et al., 2012; Ramesh and Woolford, 2016). For two ESs, ES9S on the 40S shown here (see also (Leppek et al., 2020)), as well as ES27L on the 60S previously shown by our lab (Fujii et al., 2018), humanizing ES9S or partial deletion of ES27L, respectively, result in slightly slower growth, but are sufficient to generate functional ribosomes and viable yeast cells. However, for many ESs, their complete deletion greatly reduces the level of the mutant rRNA due to biogenesis defects (Ramesh and Woolford, 2016). Thus, the length of the exchanged sequence may be crucial. Incorporating longer replacement ES sequences may be challenging if it leads to ribosome biogenesis defects extreme enough to cause lethality. In addition, if the structure of the replacement ES is a main feature important for the function addressed, the chimeric rRNA harboring the exogenous ES sequence may possibly fold into an unexpected shape since rRNA from all species are typically very structured. In this case, careful modeling or experimental probing of the structure of the ES region in question within the context of the chimeric ribosome will be important. In summary, to ensure that the ES-chimeric ribosome is designed to serve as a biologically relevant gain-of-function readout for RNA- or protein-related questions, different ES-editing sites should be specifically tested for transplantation as we have described for ES9S.

We have combined VELCRO with an *in vitro* pulldown of mRNAs to discover novel ES- mRNA interactions genome-wide. We observed a strong enrichment with hES9S and can also spatially resolve which local regions of an mRNA interact with the humanized ES. However, our design assumes that the interacting mRNA element can be reconstituted into its functional conformation in the fragmented mRNA pool. The ES9S-*Hoxa9* mRNA binding event occurs through an 18 nt-long sequence motif in the short *a9* P4 stem-loop with ES9S which to date is the shortest sequence harboring ribosome-binding activity by itself. The ES9S-interacting mRNA fragments identified by VELCRO-IP likely reveal the highest-affinity mRNA interactions with an ES as captured binding events are based on affinity. However, one cannot exclude that additional ES-binding transcripts may rely on more elaborate structures or co-factors only present within an *in vivo* setting. VELCRO-IP is thus not sensitive to potential interactions that may require other cellular components such as adaptor protein or RNA *trans*-acting factors, possible differential cellular RNA folding, or long-range interactions. We also decided for our approach and against *in vivo* RNA-RNA crosslinking methods using psoralen derivatives because psoralen derivatives only capture interactions with trans-pyrimidine configurations, which restrict sensitivity for specific interactions of mRNAs. This is especially problematic given the high GC content of many ESs. Psoralen crosslinking is furthermore hardly reversible and inefficient for lowly abundant species like mRNAs (Lipson and Hearst, 1988). Thus, this alternative crosslinking approach is impractical to be generally applicable for most potential rRNA-mRNA interactions. The strength of our method lies in its general applicability as well as in its highly specific enrichment readout. It is expected to recover the strongest RNA-RNA interactors, possibly at the expense of missing potential interactions that will not occur with *in vitro* folded mRNA fragments due to absence of additional intracellular components.

Together, VELCRO-IP allows the study of the previously uncharacterized tentacle-like rRNA expansions of the ribosome that may shape evolutionary diversity and endow greater modularity to this ancient molecular machine by mRNA-specific binding to diversify the expression of the translatome in a gene and species-specific manner.

## AUTHOR CONTRIBUTIONS

M.B., K.L., and G.W.B. conceived, and M.B. supervised the project; K.L., G.W.B. and M.B. designed the experiments, and K.L. performed experiments; G.W.B. performed RNA sequencing data analysis and GO term and statistical analysis; K.F. established the strategy for rRNA engineering and generated RPS2-Flag yeast strains; K.L. performed the rest of the experiments. M.B., K.L. and G.W.B. wrote the manuscript with input from all the authors.

## ACKNOWLEDGEMENTS

We would like to thank the Barna lab members for support and constructive criticism of the work, and for critical reading and helpful comments on the manuscript, and Gerald C. Tiu for naming VELCRO-IP. We are grateful to Katsura Asano (Kansas State University, KS, USA) and Makoto Kitabatake (Kyoto University, Japan) for providing the rDNA deletion yeast strains and the rDNA plasmids; Mary Ann Handel (The Jackson Laboratory, Bar Harbor, ME, USA) for the RPL10A antibody; and Georg Stoecklin (DKFZ-ZMBH Alliance, Heidelberg University and Mannheim University, Germany), and Davide Ruggero (both UCSF, San Francisco, CA, USA) for kindly sharing plasmids. We would like to thank John Coller and Dhananjay Wagh of the Stanford Functional Genomics Facility (SFGF) for support. This work was supported by the New York Stem Cell Foundation (M.B.), Alfred P. Sloan Research Fellowship (M.B), Mallinckrodt Foundation Award (M.B.), Pew Scholars Award (M.B.), NIH grant R01HD086634 (M.B.), a Benchmark Stanford Graduate Fellowship (G.W.B), and a Human Frontier Science Program Fellowship (K.F.). K.L. is supported by an EMBO Long-Term Fellowship (ALTF 539-2015), is the Layton Family Fellow of the Damon Runyon Cancer Research Foundation (DRG-2237-15), and is supported by the Katharine McCormick Advanced Postdoctoral Scholar Fellowship to Support Women in Academic Medicine (2019). M.B. is a New York Stem Cell Foundation Robertson Investigator.

## METHODS

### Contact for Reagent and Resource Sharing

Further information and requests for resources and reagents should be directed to and will be fulfilled by the Lead Contact, Maria Barna.

### Cell Culture and Transfection

C3H/10T1/2 (ATCC: CCL-226) cells were cultured in Dulbecco’s Modified Eagle’s Medium (DMEM, Gibco, 11965–118) containing 2 mM L-glutamine, supplemented with 10% fetal calf serum (EMD Millipore, TMS-013-B), 100 U/ml penicillin and 0.1 mg/ml streptomycin (EmbryoMax ES Cell Qualified Penicillin-Streptomycin Solution 100X; EMD Millipore, TMS-AB2-C or Gibco, 15140–122) at 37°C in 5% CO_2-_-buffered incubators. ∼0.6 × 10^6^ C3H/10T1/2 cells were seeded per well in 12-well dishes and transfected the following day with 0.8-1.6 µg of plasmid using 4 µL Lipofectamine 2000 (Invitrogen, 11668-019) and Opti-MEM (Gibco, 11058-021) according to the manufacturer’s instructions in serum-free and antibiotic-free DMEM. The medium was changed to regular DMEM 4-6 hours after transfection and cells were collected 24 hours post-transfection.

### Mice

Mice were housed under a 12 h light/dark cycle with free access to food and water. FVB/NJ (Stock# 001800) mice were purchased from the Jackson Laboratory (Bar Harbor, ME, USA) and used as wildtype. Pregnant FVB females, 3-8 months of age, were euthanized at E11.5, the uterus was dissected and embryos were taken out and placed into 1x PBS (Gibco, 14190-250). Embryos were individually collected in either TRIzol (Invitrogen, 15596) and lysed by pipetting for total RNA isolation or collected in 2 ml safe-lock tubes (Eppendorf) in 1x PBS, supernatant was removed and embryos were snap frozen in liquid nitrogen. For lysates, embryo pellets were homogenized by cryo-milling after addition of a 2.5 or 5 mm steel bead using a tissue lyser (QIAgen TissueLyser II) at 25 Hz for 15 seconds 3-6 times, and the powder was either processed directly or snap frozen in liquid nitrogen and stored at −80°C. All animal work was performed in accordance with protocols approved by Stanford University’s Administrative Panel on Laboratory Animal Care.

### Yeast Strains and Transformation

Yeast plasmids and strains (*Saccharomyces cerevisiae*) used in this paper are listed in **Table S1** and **Table S2**, respectively. Yeast strains were grown in YPD medium (10 g/L yeast extract, 20 g/L peptone, and 20 g/L glucose), YPAD medium (10 g/L yeast extract, 20 g/L peptone, 40 mg/L adenine sulfate, and 20 g/L glucose), or Synthetic Dextrose (SD) medium (6.7 g/L yeast nitrogen base, 20 g/L glucose, 1.6 g/L amino acids drop out mix (Complete Supplement Mixture, CSM, Sunrise Science Products)). All yeast strains were cultured at 30°C, unless specified otherwise. Cells were harvested in mid-log phase growth (OD_600_ = ∼0.8). Plasmid transformation of yeast cells was performed using mid-log phase cells grown in YPD, YPAD, or SD medium and standard lithium acetate-mediated transformation of 1 µg DNA and selection of transformants on SD plates of appropriate amino acids drop-out for 2-3 days at 30°C was performed.

The rDNA mutant strains were produced from the genomic rDNA deletion strain (KAY488 (NOY890)) (Nemoto et al., 2010), complemented rDNA with an exogenous plasmid, pRDN-hyg (*RDNA^hyg^ URA3*) (Nemoto et al., 2010; Wai et al., 2000), which was exchanged by plasmid shuffling to pNOY373 (*RDNA LEU2*) or derivatives containing human ES9S and 18S and 25S rRNA tags. To remove the pRDN-hyg plasmid, strains were negatively selected against the *URA3* marker gene using 1 mg/mL of 5-Fluoroorotic Acid (5-FOA) (Fisher Scientific, F10501-5.0) in SD-plates, which is processed to a toxic product by the Ura3 enzyme. To monitor rRNA processing, 5’ end processing of endogenous and tagged 18S and 25S rRNA were analyzed by RT-qPCR using pre-mature rRNA-specific or total rRNA primers (Fujii et al., 2009). Total RNA was extracted according to the manufacturer’s instructions (MasterPure Yeast RNA Purification Kit, Epicentre, MPY03100). Successful plasmid shuffling was confirmed by total RNA extraction and RT-qPCR for rRNA tags, as well as by plasmid miniprep and RT-PCR specific for the ES9S region and the 18S rRNA tag.

C-terminal Flag-tagged RPS2 strains were generated in the KAY488 (NOY890) strain by transforming 1 µg of a linear DNA template with a Kanamycin resistance cassette and 40 nt of homology arms to the target site. Selection was performed on a YPAD plate containing 200 mg/L of Geneticin (Gibco, 11811-031). Subsequently, rRNA-tagged WT and hES9S strains were generated by plasmid shuffling into this strain.

### Plasmid Construction

The following plasmids have been described previously: pSP73 (p2008) and pSP73-4xS1m (p2880) (Leppek and Stoecklin, 2014) were kindly provided by Georg Stoecklin; pSP73-4xS1m(MCS) (Leppek et al., 2020); pRF (Rluc-Fluc bicistronic; Rluc, Renilla luciferase; Fluc, Firefly luciferase reporter genes, driven by the SV40 promoter) and pRF-HCV and -EMCV (Yoon et al., 2006) were kindly provided by Davide Ruggero (UCSF); pRF derivatives containing *Hox* 5’ UTR elements and pGL3-FLB-TIE-FL containing IRES-like elements (Leppek et al., 2020).

In order to generate the series of bicistronic Rluc-IRES-Fluc pRF plasmids containing candidate 5’ UTRs from VELCRO-IP RNA-seq, full 5’ UTRs for all tested 5’ UTR-candidates and controls were either amplified from cDNA derived from E11.5 mouse mRNA reverse transcribed using SuperScript III and IV (Invitrogen, 18080044, 18090010) or from synthesized concatemerized DNA oligos (Twist Bioscience, San Francisco, USA) and inserted into the EcoRI/NcoI-sites of the bicistronic pRF vector (Yoon et al., 2006) by Gibson assembly using the NEBuilder HiFi DNA Assembly Master Mix (NEB, E2621S). Sequences were based off the ENSEMBL database (Zerbino et al., 2018) and expression profiles in input RNA-seq data. Derivatives of the plasmid pSP73-4xS1m(MCS) (Leppek et al., 2020) were generated by PCR-amplifying 5’ UTR sequences from pRF plasmids using AccuPrime Pfx DNA Polymerase (Thermo, Invitrogen, 12344024). pSP73-4xS1m(MCS) and derivatives can then be linearized at the EcoRI site downstream of the 4xS1m aptamers for run-off *in vitro* transcription.

For pNOY373-18S/25S-tag, into the yeast plasmid derivatives of pNOY373, we inserted rRNA tag sequences (Leppek et al., 2020), a 16-nt tag into 18S rRNA (Beltrame et al., 1994) and a 24-nt tag into 25S rRNA (Musters et al., 1989), for RT-PCR and RT-qPCR analysis. In a second step, the yeast ES9S was exchanged for the human ES9S in pNOY373-18S/25S-tag, which were generated by overlap extension PCR and were subsequently introduced into SacII-MluI-sites of pNOY373-18S/25S-tag, respectively. A list of all plasmids and primer sequences used are provided in **Table S1** and **Table S3**, respectively. All oligonucleotides were purchased from IDT. Mutations, cloning boundaries and coding sequences in all plasmids were verified by DNA sequencing (QuintaraBio).

### Luciferase Activity Assay after Plasmid Transfection

Transiently transfected C3H/10T1/2 cells in 12-well plates were washed twice with 1x PBS (Gibco, 14190-250) and collected by trypsinization 24 hours post-transfection for luciferase activity assays. Half the cells were used for assaying luciferase activity using the Dual-Luciferase Reporter Assay System (Promega, E1980) to measure Firefly (Fluc) and Renilla (Rluc) luciferase activities, the other half was collected in TRIzol (Invitrogen, 15596) for total RNA extraction and normalization to mRNA levels by RT-qPCR (see RT-qPCR section). For luciferase assays, cells were lysed in 60 µl of 1x passive lysis buffer of the Dual-Luciferase Reporter Assay System (Promega, E1980) and directly assayed or frozen at −20°C. After thawing, cell debris and nuclei were removed by centrifugation for 1 min at 13,000 rpm. 20 µl of supernatant was assayed for luciferase activity in technical replicates by mixing with 50 µl of Dual-Luciferase Reporter Assay System substrates. Fluc and Rluc activities were measured on a GloMax-Multi (Promega) plate reader. Luciferase reporter activity is expressed as a ratio between Fluc and Rluc which was normalized to the ratio of Fluc to Rluc mRNA levels for bicistronic pRF constructs to verify the integrity of the bicistronic mRNA construct. Each experiment was performed a minimum of three independent times. Statistical analysis was performed using unpaired two-tailed Student’s t-test.

### Quantitative RT-PCR (RT-qPCR) Analysis

Cells transfected with pRF constructs were collected in 500 µL TRIzol (Invitrogen, 15596). Total RNA was isolated from the aqueous phase using RNA PureLink columns (Thermo Scientific, Ambion, 12183018) and treated with TURBO DNase (Ambion, AM2238) twice, followed by a second RNA PureLink column purification to remove plasmid DNA. For reverse transcription-quantitative PCR (RT-qPCR) analysis, cDNA was synthesized from 100-200 ng of total RNA using iScript Supermix (Bio-Rad, 1708840) containing random hexamer primers, according to the manufacturer’s instructions. PCR reactions were assembled in 384-well plates using 2.5 µL of a 1:4-1:5 dilution of a cDNA reaction, 300 nM of target-specific primer mix and the SsoAdvanced SYBR Green supermix (Bio-Rad, 1725270) in a final volume of 10 µl per well. SYBR green detection qPCR was performed on a CFX384 machine (Bio-Rad). Data was analyzed and converted to relative RNA quantity manually or using CFX manager (Bio-Rad). Gene-specific qPCR primer sequences used for detection of mRNAs and rRNAs are given in **Table S3**.

### *In vitro* RNP affinity purification via 4xS1m-aptamers

The 4xS1m-pulldown of RNP complexes was performed similar to as previously reported (Leppek and Stoecklin, 2014). RNAs were synthesized by *in vitro* transcription: RNA elements were fused to 4xS1m aptamers by cloning 5’ UTR amplicons into the BglII/EcoRV sites of pSP73-4xS1m(MCS). 4xS1m alone served as negative control RNA. Since amplification of the highly structured 4xS1m tag by PCR is problematic, linearized pSP73 plasmids served as DNA templates. Up to 20 µg template plasmid was linearized at the EcoRI-site downstream of the 4xS1m sequence in a 50 µL reaction for 6 hours or overnight, purified with the QIAquick PCR Purification Kit (QIAgen) and used as DNA templates for run-off *in vitro* transcription using MEGAscript SP6 kit (Ambion, AM1330). A 40 µl transcription reaction contained 8 µg linear DNA template, 4 mM of each NTP (Ambion), 4 µL/ 400 U MEGAscript SP6 RNA polymerase (Ambion) and 1x SP6 MEGAscript Transcription Buffer (Ambion). After incubation for 4-6 hours at 37°C, the DNA was digested by addition of 2 µL/4 U Turbo DNase (Ambion, AM2238) for 15 min at 37°C. Synthesized RNA was purified by gel filtration using pre-packed G-50 Mini Quick Spin Sephadex RNA columns (Roche, 11814427001) according to the manufacturer’s instructions, and RNA concentration and quality was determined by Nanodrop and 4% urea-PAGE, respectively. One reaction typically yielded 50-200 µg of RNA.

For all steps in the pulldown experiments, 1.5 mL DNA/RNA LoBind tubes (Eppendorf) were used to reduce unspecific binding. Per sample, 100 µl 50% slurry of Streptavidin Sepharose High Performance (GE Healthcare) beads were washed three times with 0.5-1 ml of SA-RNP lysis buffer (20 mM Tris-HCl (pH 7.5, Ambion, AM9850G, and Ambion, AM9855G), 150 mM NaCl (Ambion, AM9759), 1.5 mM MgCl_2_ (Ambion, AM9530G), 2 mM DTT (Ambion, 10197777001), and 1 tablet/10 ml Mini Complete Protease Inhibitors, EDTA-free (Sigma-Aldrich, Roche, 11836170001) in nuclease-free water (Thermo Fisher, Invitrogen, 10977023). At each step, beads were gently pelleted at 500 rpm (∼20 x g) for 1 min at 4°C. ∼30 µg of the *in vitro* transcribed 4xS1m or 5’ UTR-4xS1m RNAs per sample for pulldown from mouse or embryo powder for protein analysis or 2.5-7.5 µg of the *in vitro* transcribed RNAs per sample for pulldown of ribosomes from yeast was renatured in 50 µl SA-RNP lysis buffer by heating at 56°C for 5 min, 10 min at 37°C, and incubation at room temperature for several minutes to refold RNA structures. The RNA was added to the 100 µl SA Sepharose slurry together with 1 µl RNasin Plus RNase inhibitor (40 U/µL, Promega, N261A). 10 µl of the supernatant was saved for extraction of input RNA using TRIzol (Invitrogen, 15596), 2.5 µl of the supernatant (input RNA) was saved for urea-PAGE analysis, and 20 µL for an input protein sample. The mixture was incubated at 4°C for 2-3 hours under rotation to permit binding of the RNA to the column. Then, beads were sedimented and 2.5 µl of the supernatant (unbound RNA) was saved for urea-PAGE analysis, while the remaining supernatant was discarded. Input and unbound RNA samples were compared side by side by 4% polyacrylamide (Ambion)/0.5x TBE (Sigma)/urea (Sigma) gel electrophoresis and SYBR Gold (10,000x, Thermo Fisher, Invitrogen, S11494) staining in 0.5x TBE to assess the efficiency of RNA coupling.

For analysis of RNA-associated proteins and RNA from yeast cells, mid-log phase cells from a 1 L SD-LEU medium culture was harvested as described in the yeast section, washed once with water, and the cell pellet was split into 16 equal aliquots into 2 ml safe-lock tubes. The yeast pellets were then snap frozen in liquid nitrogen, homogenized by cryomilling after addition of a 2.5 mm steel bead using a tissue lyser (QIAgen TissueLyser II) at 25 Hz for 30 seconds 3–6 times, or until the tissue was powderized, and the powder was either processed directly or stored at −80°C. The frozen homogenate of one aliquot (∼300 mg) was solubilized by the addition of 100 µl ice-cold RNP lysis buffer per sample and allowed to thaw for 5 min at room temperature or until thawed. Cell debris was removed by centrifugation for 5 min at 17.000 x g at 4°C, resulting in a supernatant of ∼500 µl. Yeast samples were centrifuged again for 10 min at 17.000 x g at 4°C to remove remaining cell debris. The protein concentration in the extract was determined by Nanodrop to be ∼25-70 mg/ml.

Next, the extract (∼500 µl) was pre-cleared by addition of 25 µl of a 50% slurry of Avidin Agarose (Thermo Pierce) beads, 100 µl of a 50% slurry of SA Sepharose beads, and 5 µL RNasin (Promega), and tumbling for 2 hours at 4°C. Beads were collected and discarded, and the pre-cleared lysate was supplemented with 2 µl of RNasin Plus (Promega), added onto the freshly prepared, RNA-coupled SA Sepharose matrix, and incubated at 4°C for 2-3 hours under rotation to form RNP complexes. Beads were rinsed once and washed 3 times for 2-5 min with 1 ml SA-RNP wash buffer (20 mM Tris-HCl (pH 7.5), 300 mM NaCl, 5 mM MgCl_2_, 2 mM DTT, and 1 tablet/50 ml Complete Protease Inhibitors, EDTA-free (Roche) in nuclease-free water).

For RT-qPCR analysis of RNA and WB analysis of proteins from yeast cells, elution was performed as follows. After the last wash, beads were transferred to a fresh tube and resuspended in 500 µL SA-RNP lysis buffer. 250 µL were saved and used for TRIzol extraction of bound RNA according to the manufacturer’s instructions. 15 µg GlycoBlue (Ambion, LSAM9516) was added to the RNA prior to precipitation. RNA-bound proteins were eluted from the rest 250 µL of beads by addition of 2 µg RNase A (Invitrogen, AM2271, 1µg/µL) in 30 µl Low Salt Buffer and rotation for 20 min at 4°C. The RNase A eluate was recovered, supplemented with SDS sample buffer and 8 µl of the eluate was analyzed by SDS-PAGE and WB. After RNase A elution, the beads were extracted with 30 µl 2x SDS sample buffer, 10 µl of which were analyzed by SDS-PAGE and WB. The fraction loaded of input and elution samples is expressed as percentage of the original lysate volume. For qualitative assessment of binding and elution efficiencies, an RNA fraction at each step was analysed by 4% polyacrylamide/0.5x TBE/urea gel electrophoresis and SYBR Gold staining. For qPCR analysis following RNA-IP, a fixed volume of 1:100 diluted RNA extracted from IP and input samples was used for RT. Each sample was normalized to the 18S-tag Ct values for that respective sample to control for ribosome-IP efficiency.

### Western Blot Analysis and Antibodies

Proteins were resolved on 4-20% polyacrylamide gradient Tris-glycine gels SDS-PAGE gels (Biorad, 567-1095, 456-1096) and transferred onto 0.2 µm pore size PVDF membranes (Biorad) using the semi-dry Trans-Blot Turbo system (Biorad, 170-4273). Membranes were then blocked in 1x PBS-0.1% Tween-20 containing 5% non-fat milk powder for 1 hour, incubated with antibodies diluted in the same solution for 1 hour at RT or overnight at 4°C, and washed four times for 5 min in 1x PBS-0.1% Tween-20, incubated with secondary antibodies for 1 hour in 1x PBS- 0.1% Tween-20 and washed four times for 15 min in 1x PBS-0.1% Tween-20. Horseradish peroxidase-coupled secondary antibodies (anti-mouse and anti-rabbit, GE Healthcare; anti-rat, Jackson Immunoresearch) in combination with Clarity Western ECL Substrate (Biorad, 170-5061) and imaging on a ChemiDoc MP (Biorad, 17001402) were used for detection. Antibodies were diluted in 1x PBS-0.1% Tween-20 at 1:1000 dilution either in 5% BSA (w/v) or 5% non-fat milk. The following primary antibodies were used for Western blot analysis: mouse monoclonal anti-Flag (Sigma-Aldrich, M2, F3165), anti-PGK1 (Thermo-Fisher, Novex, 459250); rabbit polyclonal anti-RPL10A/uL1 (yeast: Santa Cruz, sc-100827), anti-RPS5/uS7 (Abcam, ab58345). Rabbit polyclonal anti-RPL10A antibody was kindly provided by Mary Ann Handel (The Jackson Laboratory, Bar Harbor, ME, USA).

### Sucrose Gradient Fractionation Analysis in Yeast

For sucrose gradient fractionation of yeast cell lysates, the protocol as in (Jan et al., 2014) was used with the following adjustments. Stationary yeast cultures of cell expressing WT or hES9S rRNA in the NOY890-WT or NOY890-RPS2-Flag background were diluted to OD_600_ = 0.05 in 250 mL SD-LEU drop-out media and grown at 30°C. At mid-log phase (OD_600_ = 0.5-0.8), Cycloheximide (CHX) (Sigma Aldrich, C7698-1G) at 100 µg/ml was added into the medium and the culture was incubated for 5 min at 30°C shaking, prior to harvest omitting a water wash. Pellets were snap frozen in liquid nitrogen in 2 mL tubes. A cell pellet of a 250 mL culture was used per polysome gradient. Cell pellets were powderized by cryomilling after addition of a 2.5 mm steel bead using a tissue lyser (QIAgen TissueLyser II) 3 times at 25 Hz for 30 seconds, and the powder was processed directly. Frozen cell powder of a 250 mL culture was solubilized with 200 µL polysome lysis buffer (20 mM Tris-HCl pH 8.0 (Ambion, AM9855G), 140 mM KCl (Ambion, AM9640G), 1.5 mM MgCl_2_ (Ambion, AM9530G), 1 mM DTT (Ambion, 10197777001), 8% glycerol (Sigma-Aldrich, G5516), 1% Triton X-100 (Sigma-Aldrich, T8787), 100 µg/ml CHX (Sigma-Aldrich, C7698-1G), 100 U/ml SUPERase In RNase Inhibitor (Ambion, AM2694), 25 U/ml TurboDNase (Ambion, AM2238), Complete Protease Inhibitor EDTA-free (Sigma-Aldrich, Roche, 11836170001) in nuclease-free water (Thermo Fisher, Invitrogen, 10977023)) and vortexed. After lysis for 30 min on a rotator at 4°C, nuclei and cell debris were removed by two consecutive centrifugations (5,000 g, 5 min at 4°C, followed by 10,000 rpm, 10 min, at 4°C). Total RNA concentrations in cleared lysates were measured using a Nanodrop UV spectrophotometer (Thermo Fisher) and RNA-normalized amounts of lysates in 250 µL volume were layered onto a linear sucrose gradient (10%–45% sucrose (Fisher Scientific, S5-12) (w/v), 20 mM Tris-HCl, pH 8.0, 140 mM KCl, 5 mM MgCl_2_, 0.5 mM DTT, 100 µg/ml CHX) in nuclease-free water and centrifuged in a SW41Ti rotor (Beckman Coulter) for 2.5 hours at 40,000 rpm at 4°C. Typically, 750-1000 µg RNA was used for each sucrose gradient fractionation experiment. Fractions were collected by the Density Gradient Fraction System (Brandel, BR-188) with continuous A_260_ measurement. After collection of polysome fractions in 2 ml safe-lock tubes (Eppendorf), all fractions were individually precipitated using the Proteoextract Protein Precipitation Kit (EMD Milipore, Calbiochem, 539180-1KIT). For each 600 µL fraction, 450 µL precipitant 1 was added and incubated at −20°C for at least 1-3 hours. 10% of precipitated fractions were resolved in 26-well, 4%–20% SDS-PAGE gels (Biorad, 567-1095, 456-1096).

### VELCRO-IP RNA-seq

The Flag-pulldown of ribosome-mRNA complexes was performed analogously to as 4xS1m- mediated pulldowns from yeast, stated above. To enrich 40S ribosomal subunits, NOY890 strains that contain endogenously Flag-tagged RPS2/uS5 at the C-terminus were subjected to plasmid shuffling, as described in the yeast section, to generate tagged WT and hES9S rRNA expressing cells. Two individually isolated clones were used per strain. Cells of a 500 mL culture in SD-LEU medium were harvested when they reached mid-log phase (OD_600_ = ∼0.8-1.0). 2x 250 mL pellets were washed once with water, cells were collected in a 1.5 mL tube and flash frozen in liquid nitrogen. For lysate preparation, 250 mL pellets were powderized in liquid nitrogen using a mortar and pestle and stored at −80°C.

For the proof-of-principle pulldown experiment using 475-510 nt long *in vitro* transcripts of native, P4-native or M5-native RNAs flanked by TIE and Fluc sequences, DNA templates were amplified from monocistronic pGL3 plasmids using a SP6-flanked forward primer and Fluc-specific reverse primer (KL414/KL415) and the MEGAscript SP6 kit (Ambion, AM1330), as described in the 4xS1m pulldown section. RNA yields of 250 µg were obtained, quality was assessed by native 4-20% TBE PAGE and by SYBR Gold staining. For the Flag-pulldown experiments as described in more detail below, 5 or 7.5 µg aliquots of each *in vitro* transcript was refolded in 100 µL lysis buffer (20 mM Tris-HCl (pH 7.5, Ambion, AM9850G, and Ambion, AM9855G), 150 mM NaCl (Ambion, AM9759), 1.5 mM MgCl_2_ (Ambion, AM9530G), 2 mM DTT (Ambion, 10197777001), and 1 tablet/10 ml Mini Complete Protease Inhibitors, EDTA-free (Sigma-Aldrich, Roche, 11836170001 in nuclease-free water), and added to 50-100 µL ribosome-coupled anti-Flag M2 agarose beads and 1 µL RNasin (Promega) per reaction. Samples were rotated for 2 hours at 4°C, rinsed once and washed 3 times with 500 µL-1 ml wash buffer (20 mM Tris-HCl (pH 7.5), 300 mM NaCl, 5 mM MgCl_2_, 2 mM DTT and 1 tablet/50 ml Complete Protease Inhibitors, EDTA-free (Roche) in nuclease-free water) with rotation, before competitive 3xFlag peptide elution in 150 µL lysis buffer for 1 hour at 4°C with rotation, as stated below. 5% of the elution was used for protein analysis by WB, and 95% was subjected to TRIzol total RNA extraction and RT-qPCR analysis.

In order to generate a pool of endogenous mouse embryo mRNAs as RNA input for the ribosome-IP, up to 10 stage 11.5 embryos per FVB female were harvested as described in the mouse section, individually collected in 2 ml Eppendorf tubes, washed once with 1x PBS (Gibco, 14190-250), and lyzed in 1 mL TRIzol (Invitrogen, 15596) by pipetting and vortexing, and addition of another 800 µL TRIzol. Embryo lysates were stored at −80°C until total RNA extraction. From each embryo, 150-200 µg total RNA was obtained. From total RNA, poly(A) mRNA was isolated on oligo(dT) beads using the Oligotex mRNA Mini Kit (QIAgen, 70022) or Poly(A) Purist MAG kit (Invitrogen, AM1922) according to the manufacturer’s instructions, which yielded ∼5 µg mRNA (2-3%) of 150-200 µg total RNA per embryo. Purified embryo mRNA was fragmented to 100-200 nt RNA fragments by magnesium-buffer based degradation using the NEBNext Magnesium RNA Fragmentation Module (NEB, E6150S). Fragmentation was optimized for time and RNA input amount monitoring RNA size using the mRNA Pico Chip (Agilent) on a Bioanalyzer (Agilent), and by 8% denaturing urea-PAGE and SYBR Gold staining. mRNA fragmentation was initially optimized using mRNA isolated from mouse C3H10T1/2 mesenchymal cells instead of embryo tissue and the yield of purified mRNA was identical from different source material. We tested input mRNA amounts of 250 ng, 500 ng and 1 µg mRNA over a time course of 0-10 min, since the manufacturer’s protocol only indicated use for up to 250 ng mRNA. Fragmentation of 1 µg mRNA aliquots for 5 min at 94°C in 1x Fragmentation Buffer (NEB) was optimal to obtain a pool of 100-200 nt fragments. Reactions were quenched on ice and by addition of 1x Stop Solution (NEB). Immediate isopropanol-based precipitation recovered 75- 95% of input mRNA as mRNA fragments in water.

For Flag-pulldown of Flag-tagged yeast 40S, powderized yeast lysates of a 250 mL culture per three samples were dissolved in 500 µL lysis buffer and the tube was washed with another 200 µL lysis buffer. Lysates were cleared by centrifugation for 5 min at 17,000 g at 4°C and 2 min at 17,000 g at 4°C, and 800 µL lysate was recovered. RPS2-Flag tagged 40S ribosomes were immuno-precipitated by addition of 50 µL washed anti-Flag M2 affinity agarose gel (Sigma Aldrich, A2220-5mL) and 5 µL RNasin Plus (Promega) per sample to 800 µL lysate and 1.5-2 hours of rotation at 4°C. Beads were washed 3 times with 500 µL lysis buffer and bound ribosomes were resuspended by addition of 200 µL lysis buffer. 10 µg fragmented mRNA from E11.5 FVB embryos in 40 µL per sample were pooled for 6 samples. 5 µL was saved as an input RNA sample for sequencing. Pooled mRNA was refolded in lysis buffer in a total volume of 600 µL as described in the 4xS1m pulldown section and used as input for 6 samples. 10 µg refolded RNA in 100 µL was added to 100 µL ribosome-coupled 50% beads, 3 µL RNasin (Promega) and 100 µL lysis buffer for a total volume of 300 µL for IP by rotation for 2 hours at 4°C. Bound ribosome-mRNA fragment complexes were rinsed once with 1 mL lysis buffer and washed 3 times with wash buffer for 5 min tumbling at 4°C. Samples were then eluted off the anti-Flag beads using competitive 500 µg/mL 3xFlag peptide (Sigma-Aldrich, F4799-4mg) elution in 150 µL lysis buffer by rotation for 1 hour at 4°C. 5% of the elution was used for protein analysis by WB, and 95% was subjected to TRIzol total RNA extraction and library preparation.

### Library Preparation and Deep Sequencing

5 µg total RNA isolated from Flag elution samples were treated with Yeast RiboZero Gold (Illumina, MRZY1306) according to the manufacturer’s instructions to remove yeast rRNAs from the samples. From the remaining fragmented RNA in water (10 µL, yield 80-160 ng RNA), 30 ng of elution and mRNA fragment input samples were used for library preparation. Library preparation for deep sequencing was performed using the NextFlex Rapid Directional qRNA-Seq Library Prep Kit (Perkin Elmer, Bioo Scientific, NOVA-5130-01D) according to the manufacturer’s instructions using 7 unique barcodes. In brief, the standard protocol was applied with the following changes: the initial fragmentation step was omitted and PCR amplification was performed using 16 cycles. DNA fragments were purified for Illumina sequencing, subjected to analysis using the High Sensitivity DNA Assay (Agilent) on a Bioanalyzer (Agilent) and all DNA libraries were pooled to a final concentration of 4 nM. Sequencing was performed at the Stanford Functional Genomics Facility (SFGF) at Stanford University, on the Illumina NextSeq 550 instrument, using 2x 75 nt paired-end sequencing and the following library design: AATGATACGGCGACCACCGAGATCTACACTCTTTCCCT ACACGACGCTCTTCCGATCTNNNNNNNNT-insert- NNNNNNNNAGATCG GAAGAGCACACGTCTGAACTCCAGTCACBBBBBBBBATCTCG TATGCCGTCTTCTGCTTG, where N is the 2x 8 nt unique molecular index, and B is the 8 nt sample barcode.

### VELCRO-IP RNA-seq Data Analysis: Read Alignment and Quantification

First, for removal of adapter sequences, low quality bases, and short reads, we use cutadapt (Martin, 2011) to trim Illumina adapter sequences and <Q20 bases. Reads <40 nt were removed. Parameters: cutadapt -m 40 -a AGATCGGAAGAGCACACGTCTGAACTCCAGTCAC -A AGATCGGAAGAGCGTCGTGTAGGGAAAGAGTGTAGATCTCGGTGGTCGCCGTATCA TT --nextseq-trim=20. Next, for UMI extraction, we used umi_tools (Smith et al., 2017) to extract the UMI region (first 8 bases). Parameters: umi_tools extract --bc-pattern=NNNNNNNN --bc- pattern2=NNNNNNNN. We additionally remove 1 base from 5’ end of the reads, which is the A/T nucleotide overhang from the ligation reaction during library preparation. For splice-aware alignment using STAR (Dobin et al., 2013), we used STAR to align the reads to a reference genome/transcriptome. STAR reference is built using a combination of yeast genome (sacCer3), mouse genome (mm10), mouse rDNA sequence (GenBank GU372691), and mouse transcript annotations (GENCODE vM18). Only uniquely mapped reads were retained. Parameters: STAR --sjdbOverhang 66 --outFilterMultimapNmax 1 --alignEndsType EndToEnd --alignIntronMax 1000000 --alignMatesGapMax 1000000 --alignIntronMin 20 --outFilterMismatchNmax 999 -- alignSJDBoverhangMin 1 --alignSJoverhangMin 8 --outFilterType BySJout. While the majority of the reads mapped to yeast mRNAs that we believe reflect background binding from the initial ribosome-IP (∼20 million reads), 1-3% mapped to mouse mRNAs which corresponds to ∼500,000 reads per sample. For deduplication using UMI, we used umi_tools to deduplicate the alignments. Deduplicated alignments are re-aligned using STAR and the same parameters as before. Parameters: umi_tools dedup --paired --buffer-whole-contig. For read quantification, we used bedtools (Quinlan and Hall, 2010) to count alignments over 200 nt sliding windows with step size of 100 nt across mouse genome.

### VELCRO-IP RNA-seq Data Analysis: Enrichment Analysis

For data matrix and normalization, each cell in the data matrix is the read count, where rows are 200 nt genomic windows and columns are the samples. We discarded rows whose sum of counts across all six mutant and wild-type samples was <30. We used the TMM (Robinson and Oshlack, 2010) method to calculate normalization factors. Counts divided by normalization factors were used for plotting tracks along the transcript. Tracks are plotted using wiggleplotR (Alasoo et al., 2015). Each genomic window is annotated as 5’ UTR, ORF, or 3’ UTR based on any overlap with any isoform present in the GENCODE vM18 annotation. For statistical significance of enriched windows, we use voom (Law et al., 2014)-limma (Ritchie et al., 2015) to model mean-variance bias and calculate moderated t-statistics and p-values for the difference in mutant versus wild-type samples. We noted the heavy tailed histogram of the t-statistics suggesting high proportion of non-null windows and used locfdr (Efron, 2010) approach to estimate local false discovery rates. All reported FDR values in the manuscript are locfdr estimates. Locfdr parameters: bre=150, df=25, pct=0, nulltype=1, type=0, mlests=(-0.5, 1.0). To test overrepresentation of enriched windows across 5’ UTR-CDS-3’ UTR regions, we performed permutation based chi-square test of independence on the contingency table of regions that the windows overlap versus whether the FDR for enrichment of windows were <=0.05. For Gene Ontology (GO) term enrichment, GO terms and gene mappings were obtained from Bioconductor annotation package org.Mm.eg.db (version 3.6.0). We used topGO (Alexa et al., 2006) to perform enrichment analysis. We choose the combination of Fisher’s exact test and weight01 algorithm for handling local similarities between GO terms. Genes that have at least one window with FDR <= 0.05 are used as the positive set. All genes that have at least one window tested are used as the background. For the reported list of GO terms in the manuscript, the following criteria are true: observed/expected ratio >= 2, minimum number of observed genes >= 3, Fisher’s exact test FDR<=0.05, and weight01- conditioned Fisher’s exact test p-value <= 0.05. FDR for Fisher’s exact test is estimated by permutation of the gene labels of the positive set.

### Data Sources

For the multiple sequence alignment (MSA) and conservation analysis of ES9S and surrounding 18S rRNA sequence, the following GenBank 18S rRNA sequences were retrieved for eukaryotic species from the NCBI database as data sources and references and aligned by Multiple Alignment using Fast Fourier Transform (MAFFT, MView, EMBL-EBI webtools) with default settings: mouse (*Mus musculus*; accession number NR_003278.3), human (*Homo sapiens*; M10098.1), chicken (*Gallus gallus*; AF173612.1), African clawed frog (*Xenopus laevis*; X02995.1), zebrafish (*Danio rerio*; NR_145818.1); juvenile axolotl (*Ambystoma mexicanum*); and yeast (*Saccharomyces cerevisiae*; J01353.1).

### Quantification and Statistical Procedures

In all figures, data was presented as mean, SD or SEM as stated in the figure legends, and *p ≤ 0.05 was considered significant (ns: p > 0.05; *p ≤ 0.05; **p ≤ 0.01; ***p ≤ 0.001; ****p ≤ 0.0001). Blinding and randomization were not used in any of the experiments. Number of independent biological replicates used for the experiments are listed in the figure legends. Tests, two-tailed unpaired Student’s t-test if not stated otherwise, and specific p-values used are indicated in the figure legends. In all cases, multiple independent experiments were performed on different days to verify the reproducibility of experimental findings. For mouse experiments, embryos from multiple litters were used to avoid litter-specific bias.

## ACCESSION NUMBERS

RNA sequencing data from VELCRO-IP RNA-seq experiments were deposited in the Gene Expression Omnibus (GEO) under accession number GSE141382.

## SUPPLEMENTAL INFORMATION

### SUPPLEMENTAL FIGURE LEGENDS

**Figure S1.**
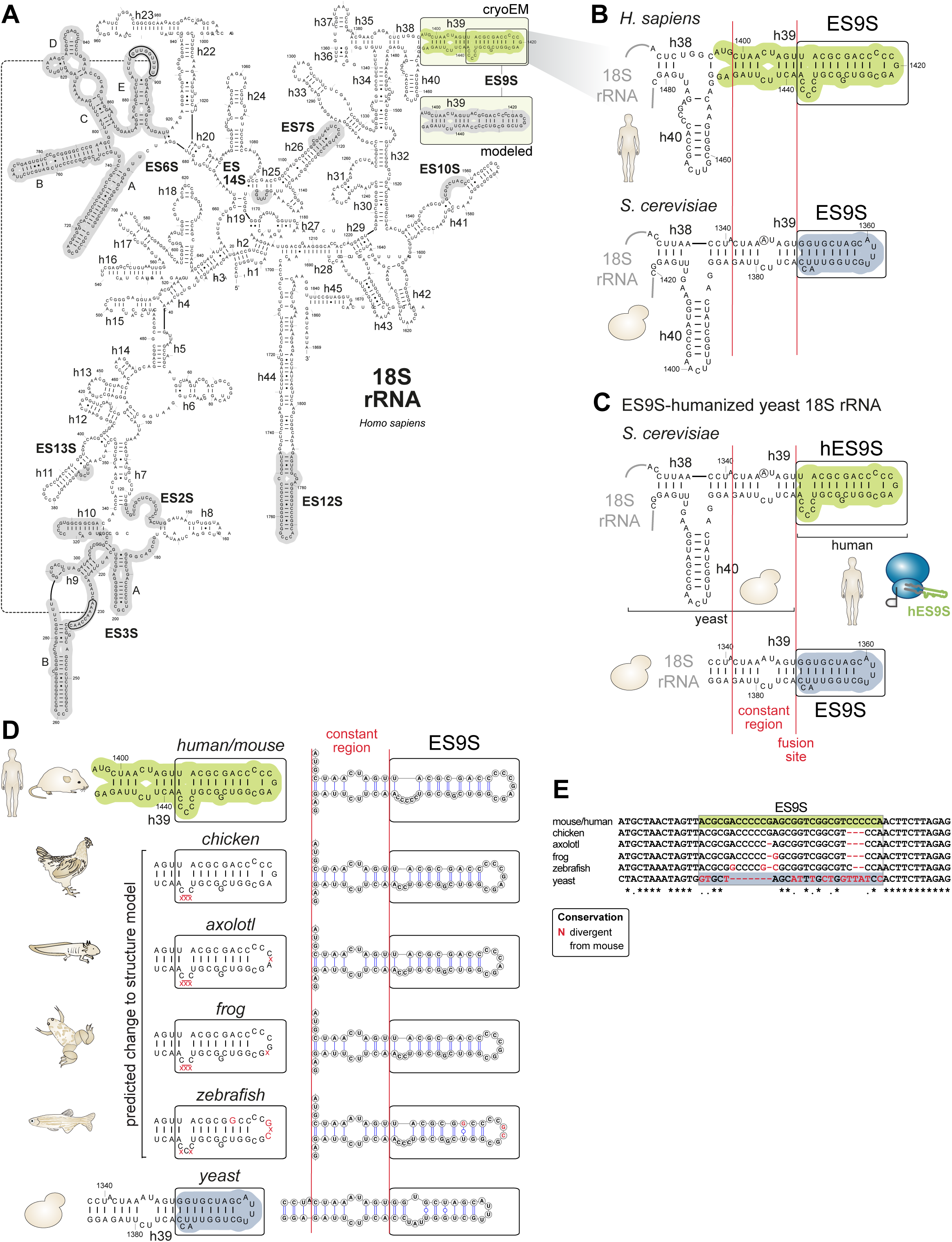
Confirmation of interspecies sequence variation of ES9S rRNA region. Related to Figure 1. (A) Secondary structure model of the human (*H. sapiens*) 18S rRNA adapted from (Anger et al., 2013). rRNA expansion segment regions are highlighted in grey. Nucleotide positions, helices and ESs are numbered. The boxed region of ES9S is shown for the model (grey) and based on our cryo-EM data (green; (Leppek et al., 2020)). (B) Secondary structure models of the human and baker’s yeast (*S. cerevisiae*) 18S rRNA region containing ES9S, highlighted in green and blue, respectively. The structure of the distal human ES9S (boxed region in A and B) was revised based on cryo-EM data (Leppek et al., 2020). (C) Secondary structure model of the result of engineered yeast 18S rRNA after exchange of the yeast ES9S with the human one (hES9S, green). Constant region (h39) and ES9S-fusion site selected for engineering chimeric 18S rRNA are indicated in red. (D) Predicted structural changes in the ES9S region of 18S rRNA with regards to species-specific variation in sequence. We annotated sequence changes and their possible effect on the ES9S structure in red that are divergent from the identical human and mouse ES9S, as derived from RT-PCR analysis of the ES9S region in 18S rRNA using cDNA generated from total RNA derived from many different species (E11.5, stage E11.5 FVB mouse embryo; chicken, *Gallus gallus*; axolotl, *Ambystoma mexicanum*; frog, X. l., *Xenopus laevis*; zebrafish, *Danio rerio*; yeast, S. c., *Saccharomyces cerevisiae*) and primers specific for the 18S rRNA region containing ES9S in the center (see Figure 1A-C, partially reproduced from Figure 1A). Secondary structure models of ES9S of different species were predicted using Vienna RNAfold (http://rna.tbi.univie.ac.at) and visualized using VARNA (http://varna.lri.fr) with default settings. (E) Alignment of RT-PCR confirmed sequences of ES9S-containing 18S rRNA-sequence used for the structure model predictions in (D). Partially reproduced from Figure 1C.

**Figure S2.**
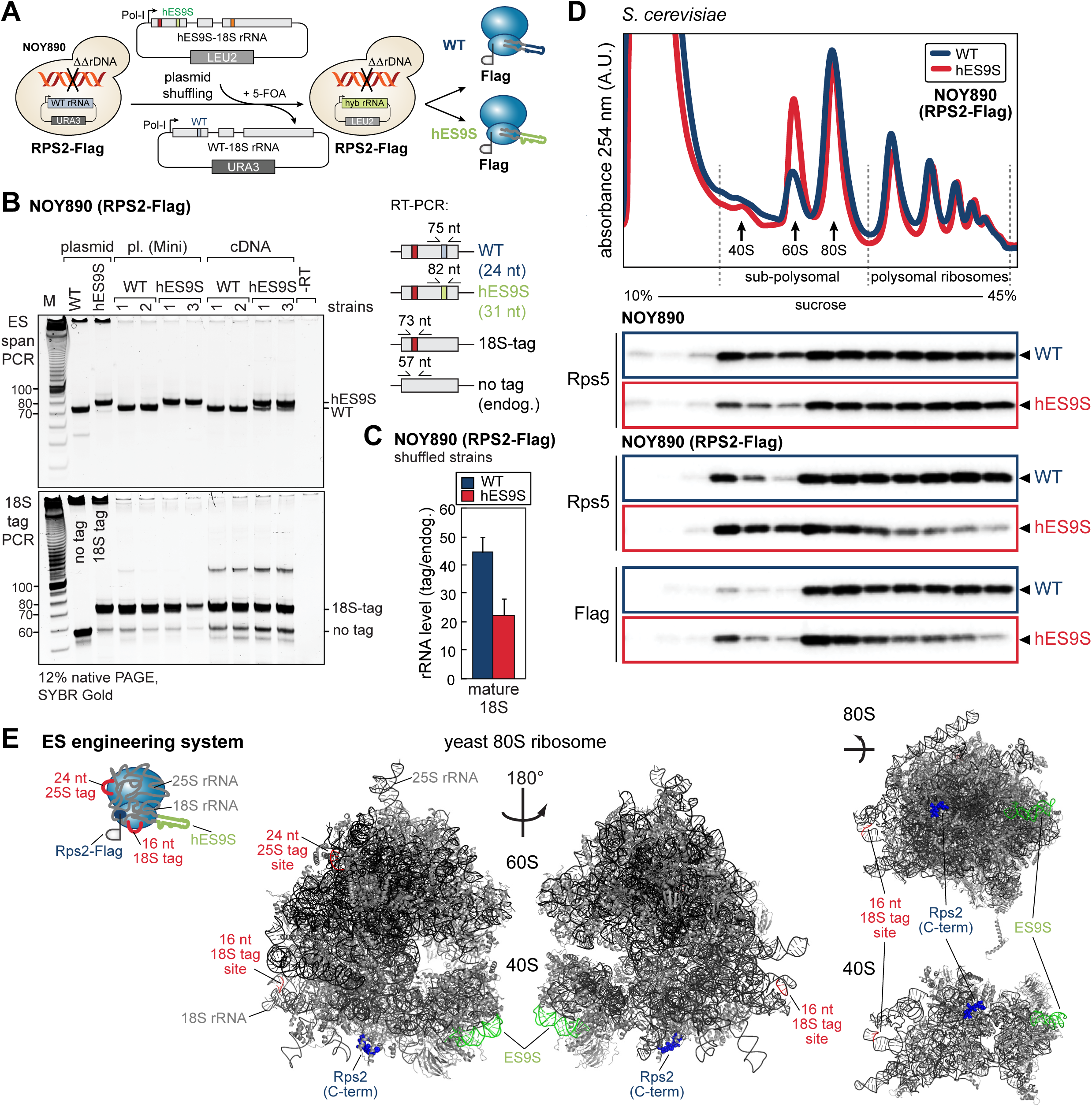
Plasmid shuffling in yeast and strain characterization. Related to Figure 1, 2. (A) A yeast strain containing the plasmid-encoded chimeric 18S rRNA is generated by plasmid shuffling. Schematic of the plasmid shuffling approach to generate yeast strains (NOY890, RPS2-Flag) that contain a homozygous knock-out of the rDNA locus (NOY890) and generate ribosomes exclusively from plasmids. All rDNA plasmids contain unique 18S and 25S rRNA sequence tags. 5-FOA-based selection of transformed yeast cells allows for isolation of clones that retain a transformed *LEU2*-plasmid (pNOY373) and lost the original *URA3*-plasmid (pNOY373). Successful plasmid exchange from *URA3* (WT) to *LEU2* (tagged WT or hES9S)-plasmids in isolates is achieved by growth on SD-*LEU2*, and SD+5-FOA but not on SD-*LEU/URA*. (B) RT-PCR analysis using ES9S-specific primers that span ES9S allow analysis of expression of WT or hES9S 18S rRNA due to a PCR product of 7 nt difference in length between WT and hES9S (ES span PCR). Similarly, the presence of the 18S tag can be distinguished from WT rRNA (18S tag PCR). Total RNA for cDNA synthesis or plasmid DNA was extracted from clones and used for RT-PCR. Plasmid-derived PCR products serve as controls. PCR products were resolved by 12% native PAGE and stained with SYBR Gold. Two independent isolates of tagged-WT and tagged- hES9S strains (NOY890/RPS2-Flag background) used in this study are presented. RT-PCR specific for the 18S rRNA tag confirms presence of the tag in transformed plasmid and derived mature 18S rRNA. A 10 bp DNA ladder (Invitrogen) was loaded as reference. (C) For yeast strain characterization after plasmid shuffling and isolation of clones, RT-qPCR analysis using specific primers for rRNA tags and endogenous rRNA is used to derive a tag/endogenous rRNA level that assesses the substitution rate of WT with tagged-WT or tagged- hES9S ribosomes present in isolated strains. For NOY890/RPS2-Flag strains, for 44 and 22 tagged WT and hES9S ribosomes, respectively, one endogenous plasmid-derived WT ribosome is left in the cell. (D) Sucrose gradient fractionation analysis of yeast lysates derived from WT and hES9S-stains in the background of NOY890 and NOY890/RPS2-Flag, containing scarless C-terminal RPS2-Flag (Jan et al., 2014), on 10-45% sucrose gradients (n = 3). In comparison to WT rRNA-containing cells, humanized ribosome-containing cells show a slight growth defect. Polysome traces demonstrate proper ribosomal assembly. Incorporation of the Flag tag into polysomes demonstrates its non-perturbative nature. (E) Mapping of the components of the ES engineering system onto the cryo-EM structure of the yeast 80S and 40S ribosome (PDB: 4V6I). The sites of rRNA tag insertion, the last 10 amino acids of the C-terminus of Rps2, and ES9S are highlighted according to the schematic representation.

**Figure S3.**
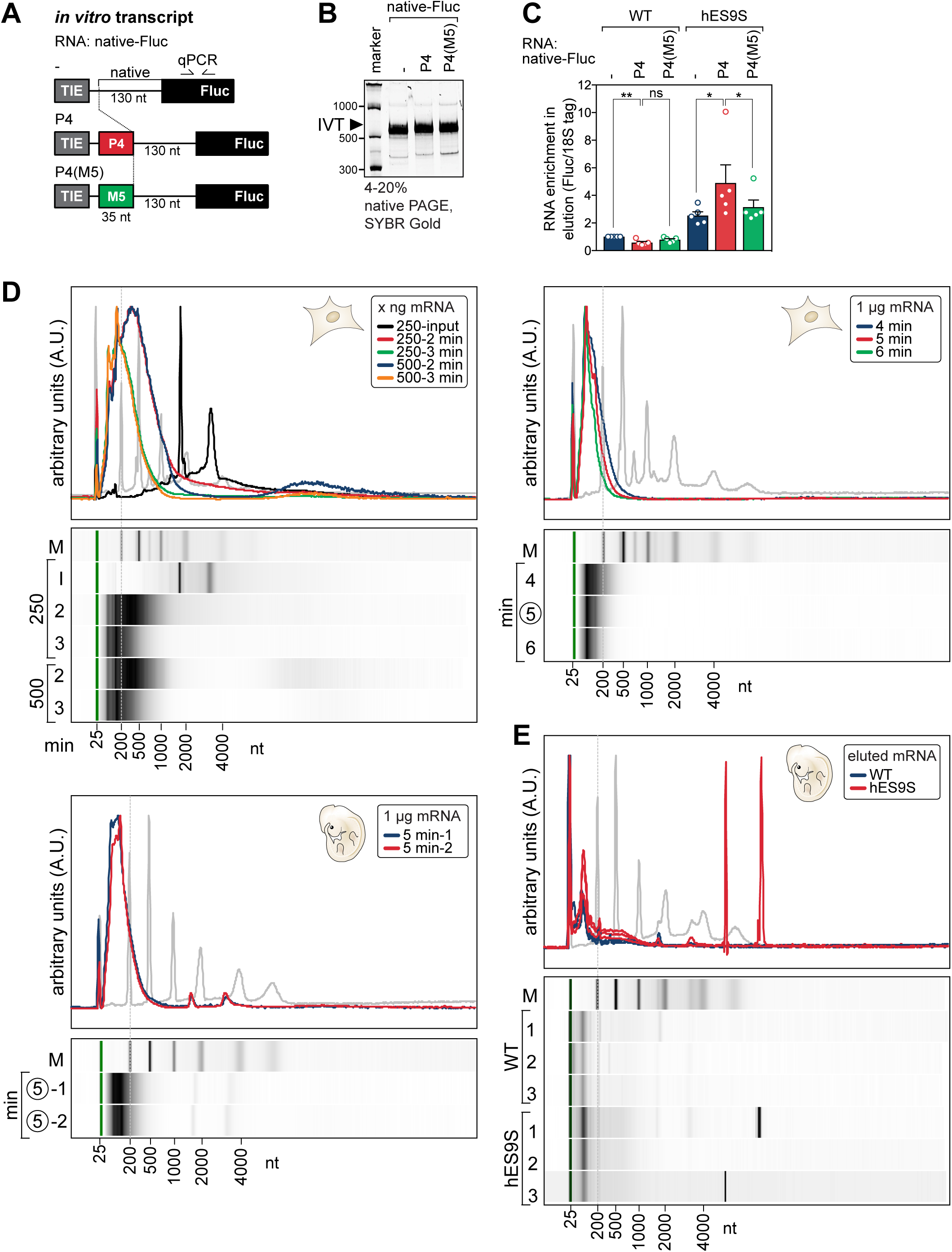
VELCRO-IP RT-qPCR serves as a proof-of-principle to identify novel hES9S-interacting 5’ UTRs and mRNA fragmentation. Related to Figure 3. (A) Schematic of *in vitro* transcripts used for the proof-of-principle experiment of the VELCRO-IP RT-qPCR. Reproduced from Figure 3B. (B) For qualitative analysis of the integrity of *in vitro* transcripts, RNAs were subjected to 4-20% polyacrylamide/TBE/native PAGE and visualized by SYBR Gold staining. (C) Analysis of total RNA in the 3xFlag peptide elution by RT-qPCR using same volumes of RNA per sample for the RT. Normalization of Ct values for Fluc to the 18S rRNA tag internally controls for ribosome-IP efficiency per sample. The native/WT sample was used to normalize for fold enrichment of RNA binding (set to 1). Representation of the raw data in Figure 3D. Average RNA fold enrichment, SEM, n = 5; ns, not significant. (D) Full view of the Bioanalyzer (Agilent) quantification and electronic gel analysis in Figure 3G, H is shown for optimization of mouse mRNA fragmentation from C3H/10T1/2 cells and stage E11.5 mouse embryos. The marker (M, grey) is overlaid for reference. (E) Full view of the Bioanalyzer (Agilent) quantification and electronic gel analysis in Figure 4B is shown for the eluted and yeast rRNA-depleted mouse embryo RNA from three independent replicates of WT and hES9S VELCRO-IP experiments. The marker (M, grey) is overlaid for reference.

**Figure S4.**
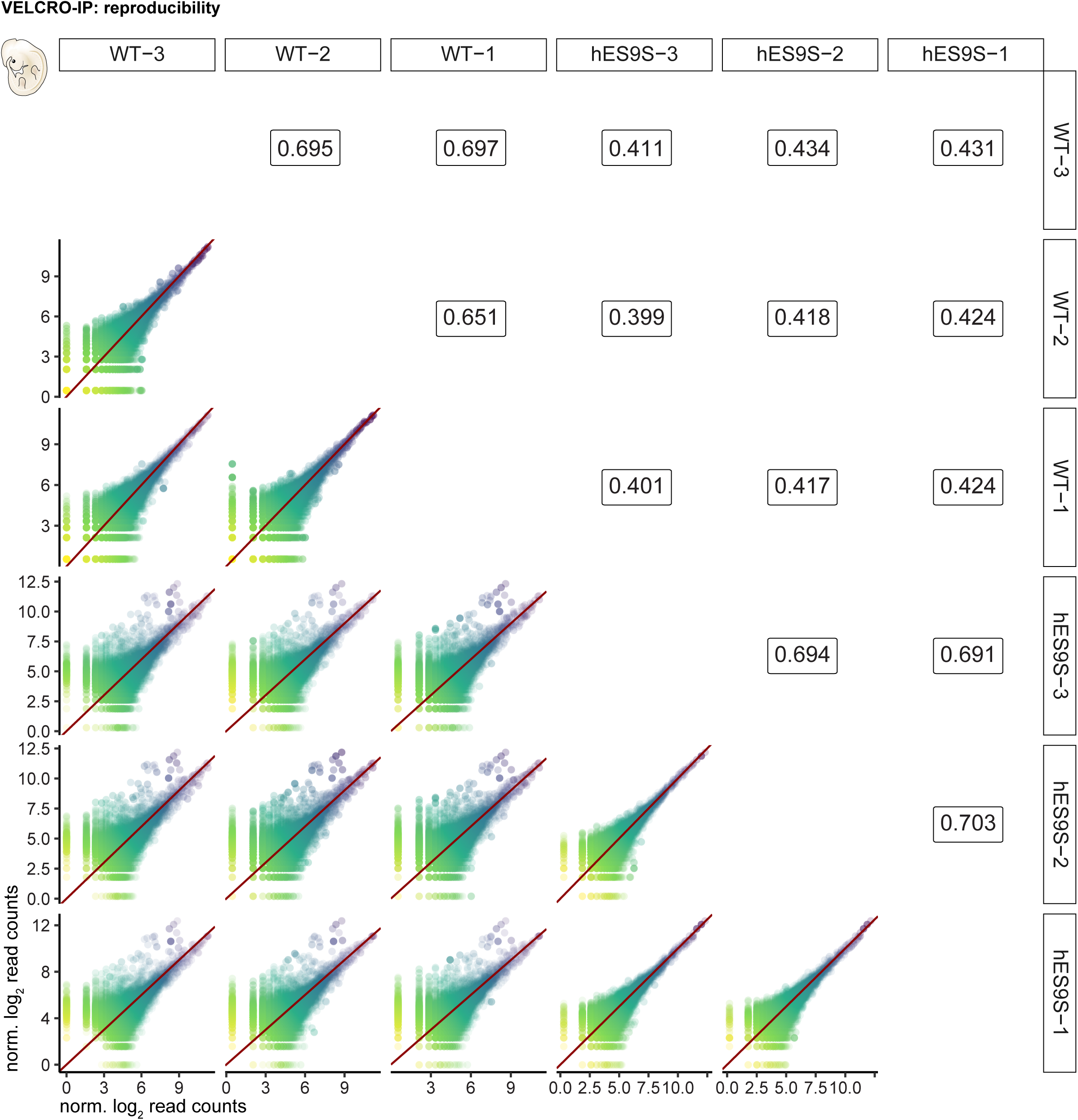
Reproducibility of VELCRO-IP RNA-seq. Related to Figure 4. A matrix comparing every possible pair of individual VELCRO-IP RNA-seq samples (three replicate samples per condition, hES9S and WT). Lower triangle: scatter plots of normalized log read counts, colored by expression level. Upper triangle: Pearson correlation coefficient.

**Figure S5.**
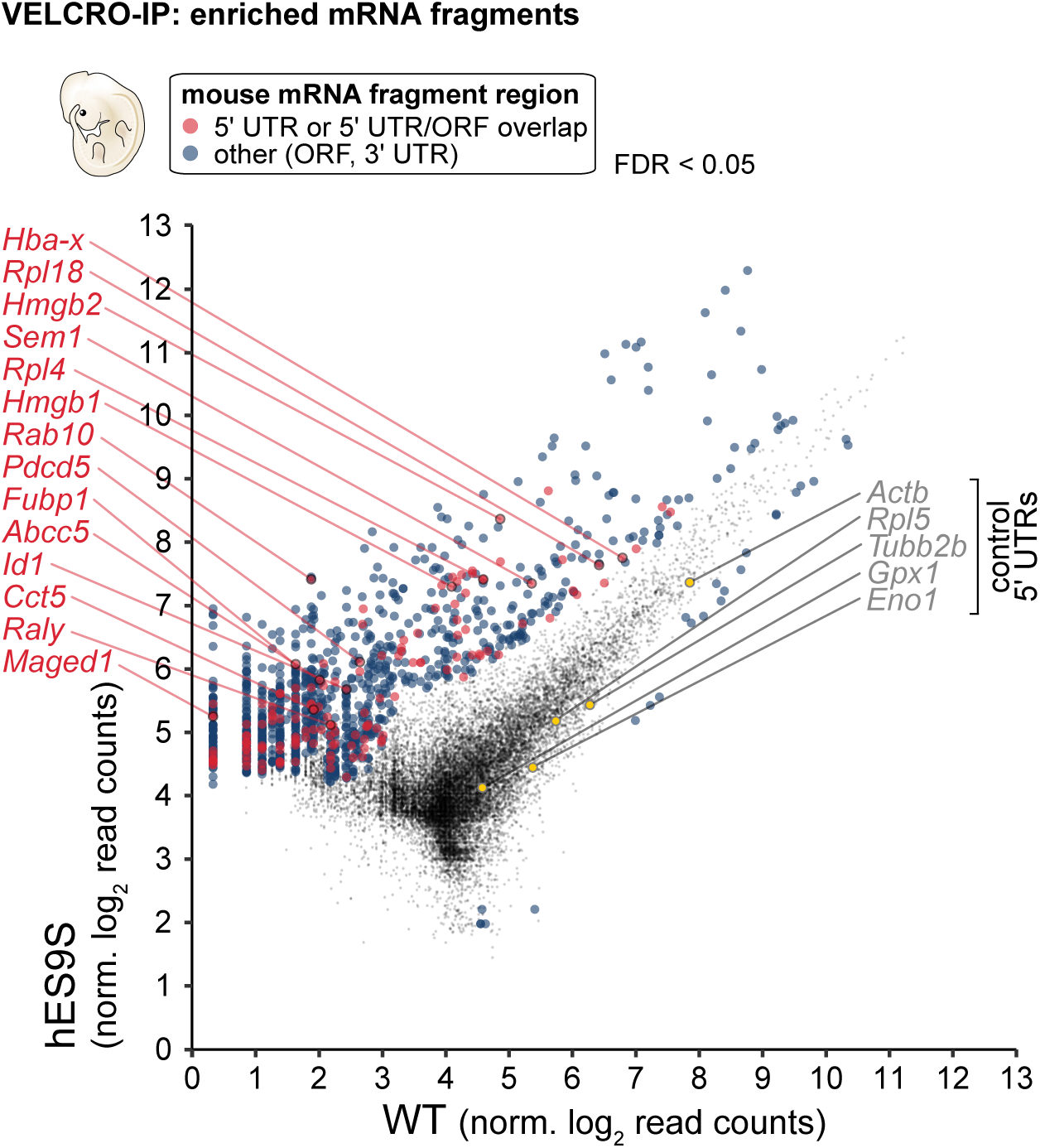
VELCRO-IP RNA-seq identifies novel hES9S-interacting 5’ UTRs. Related to Figure 4. RNA-seq results of independent replicates (n = 3) for each WT and hES9S samples. Normalized log read counts are presented for WT and hES9S-enriched mRNA fragments. Fragments less than FDR < 0.05 are colored according to the region in the mRNA they map to. Fragments mapping to 5’ UTR and overlapping 5’ UTR/ORF (red) are highlighted compared to other regions (ORF and 3’ UTR, blue). We label mouse genes for which we identified enriched fragments in the 5’ UTR and/or 5’ region of the ORF and for whose 5’ UTRs we performed validation experiments. Five control 5’ UTRs are highlighted in yellow that are equally bound to both WT and hES9S 40S subunits and served as negative controls. Corresponds to Figure 4E. See also **Table S4**.

**Figure S6.**
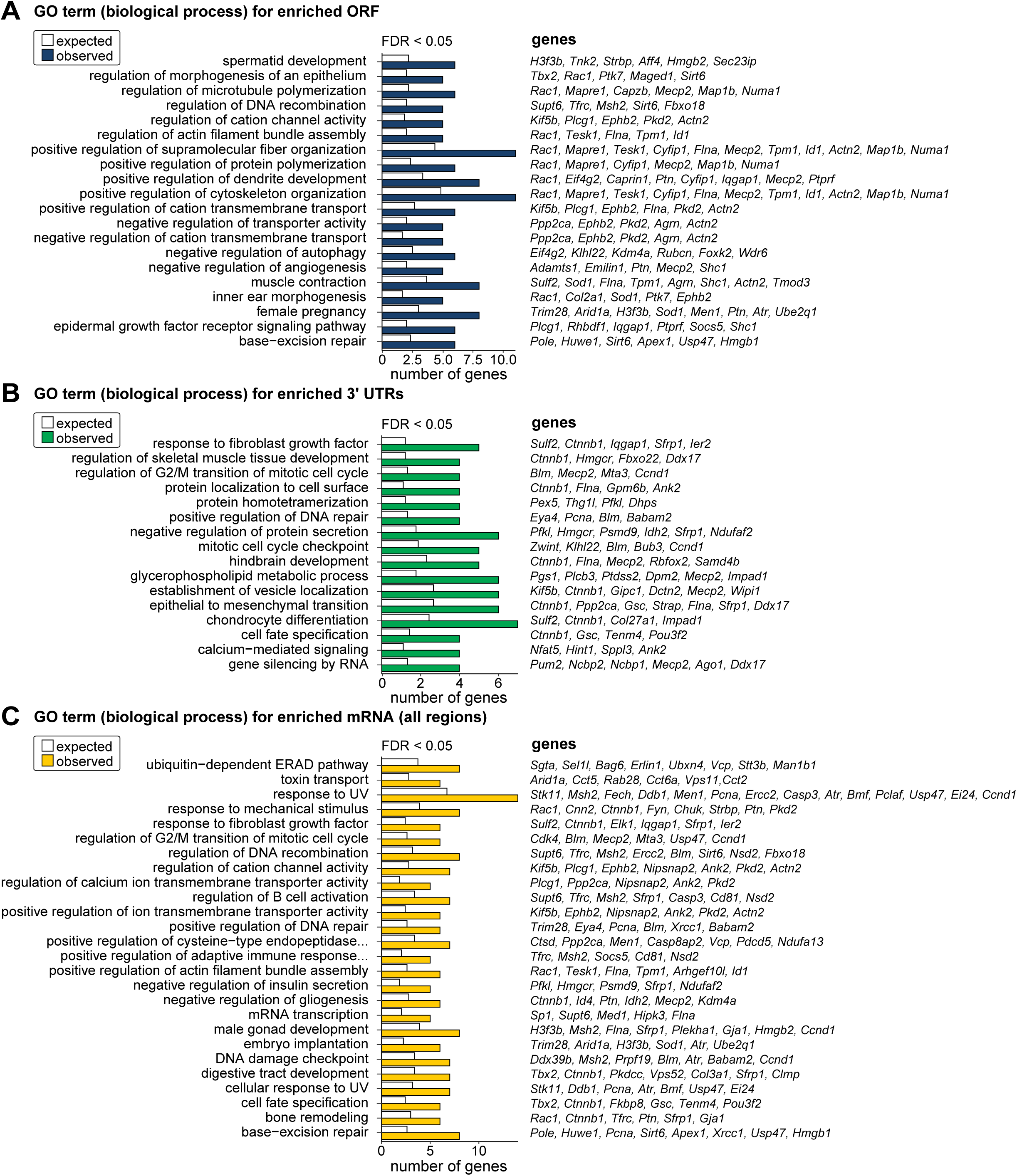
GO-terms of novel hES9S-interacting mRNA regions. Related to Figure 4. (A) GO term analysis as in Figure 4G for biological process of total ORF regions (FDR < 0.05, n = 3) enriched with hES9S. Displayed are the expected and observed frequency of genes in categories. mRNA fragments present in either WT or hES9S samples were used as the background population (expected frequency). Also see **Table S5**. (B) GO term analysis as in (A) for biological process of total 3’ UTR regions (FDR < 0.05, n = 3) enriched with hES9S. (C) GO term analysis as in (A) for biological process of the full mRNA (all regions) regions (FDR < 0.05, n = 3) enriched with hES9S.

**Figure S7.**
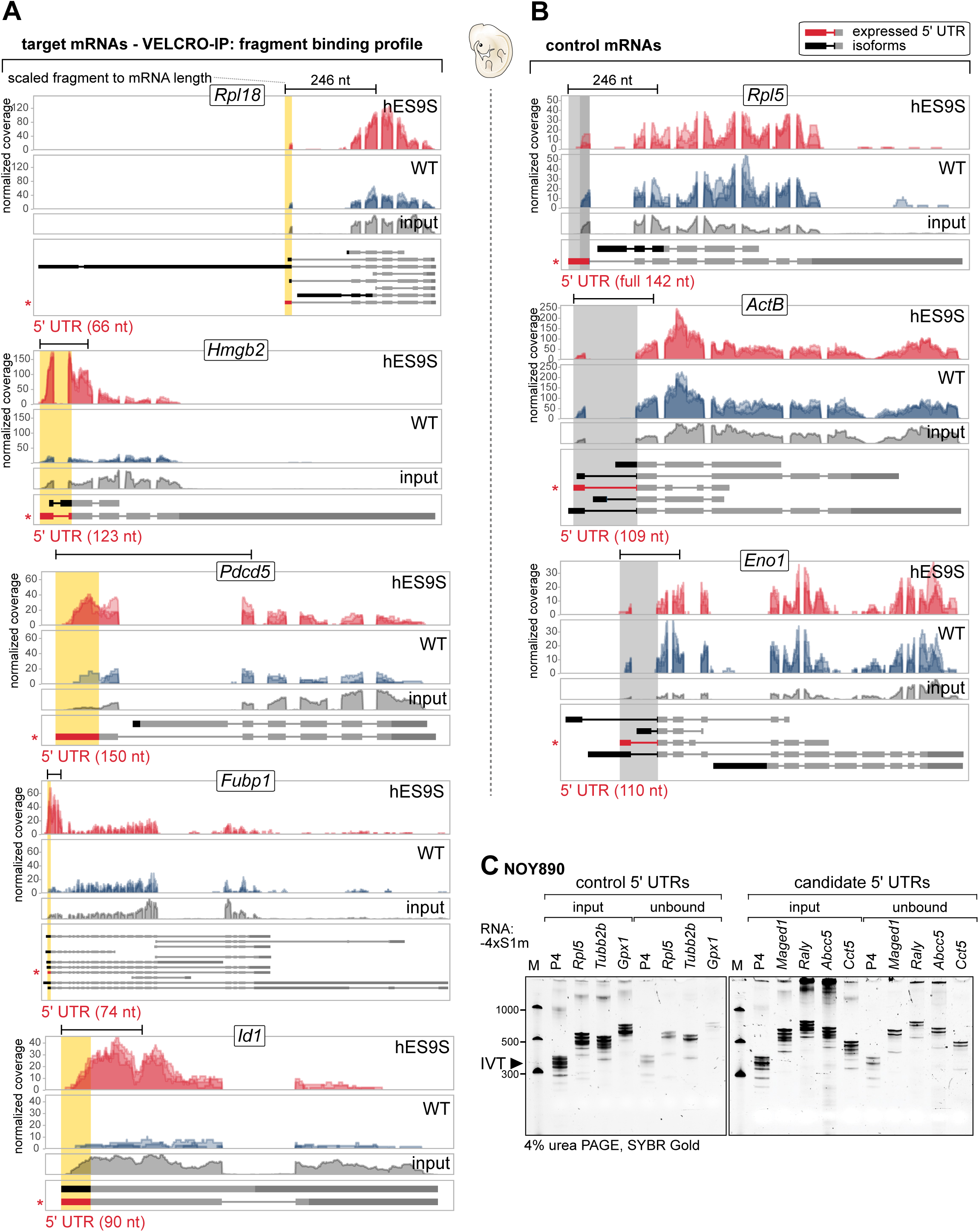
VELCRO-IP mRNA binding pattern and validation of novel hES9S- interacting 5’ UTRs. Related to Figure 5, 6. (A) mRNA binding profile as coverage plots for candidate hES9S-target genes whose 5’ UTR-overlapping windows are significantly enriched in the hES9S over WT samples (FDR < 0.05, n = 3). The other five out of the total tested 14 genes not shown in Figure 5A are given here. Normalized per base coverage of individual biological replicate libraries for WT (blue) and hES9S (red) samples is plotted (above). All mRNA isoforms annotated in ENSEMBL are displayed below. Exon lengths are to scale while intron lengths are pseudo-scaled. The read coverage of the input mRNA fragments (grey) are also plotted for reference. 5’ UTR regions for the most likely expressed mRNA isoform in embryos is highlighted in red and the corresponding regions in the tracks is shaded in yellow. The 5’ UTR region picked for further experimental validation corresponds to the asterisk-marked isoform. The mRNA fragment length for each gene is scaled according to the mRNA length for the individual genes presented. See also Figure 5A. (B) The same analysis as in (A) was performed for the other three of total five control 5’ UTRs that we found to equally bind to either WT or humanized 40S. 5’ UTR regions for the most likely expressed mRNA isoform in embryos is highlighted in red and the corresponding regions in the tracks, that also indicate no specific enrichment, is shaded in gray. Corresponds to Figure 5B. (C) A 4xS1m pulldown experiment with the focus on the comparison of full-length control and candidate hES9S-interacting 5’ UTRs for their ability to bind to tagged-WT and tagged-humanized 40S subunits was performed. *In vitro* transcribed RNAs fused to 4xS1m aptamers were coupled to SA-sepharose beads for 4xS1m pulldown using WT and hES9S ribosome expressing yeast strains to generate cellular extracts as input. Coupled beads were incubated with cell extracts, washed and eluted using RNase A to release RNA-bound proteins. Input and unbound samples were taken before and after incubation of RNAs with beads. To monitor coupling efficiency, 10% of the input and unbound RNA fraction of each sample was resolved by 4% denaturing polyacrylamide/TBE/urea PAGE and visualized by SYBR Gold. Representative of n = 3 is shown. Low Range ssRNA Ladder (NEB) was loaded for reference. Corresponds to Figure 6C.

## STAR METHODS

### Key Resources Table

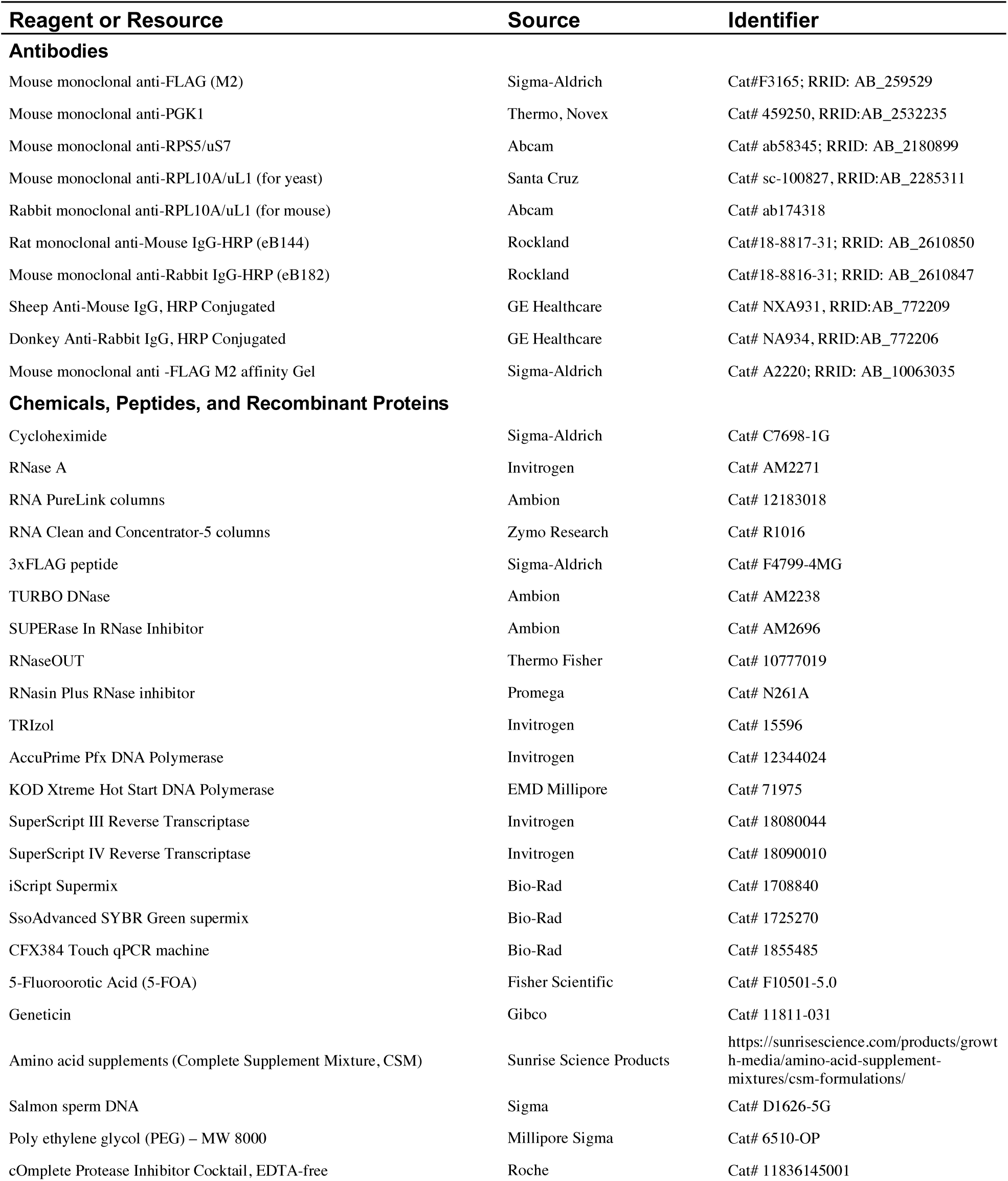
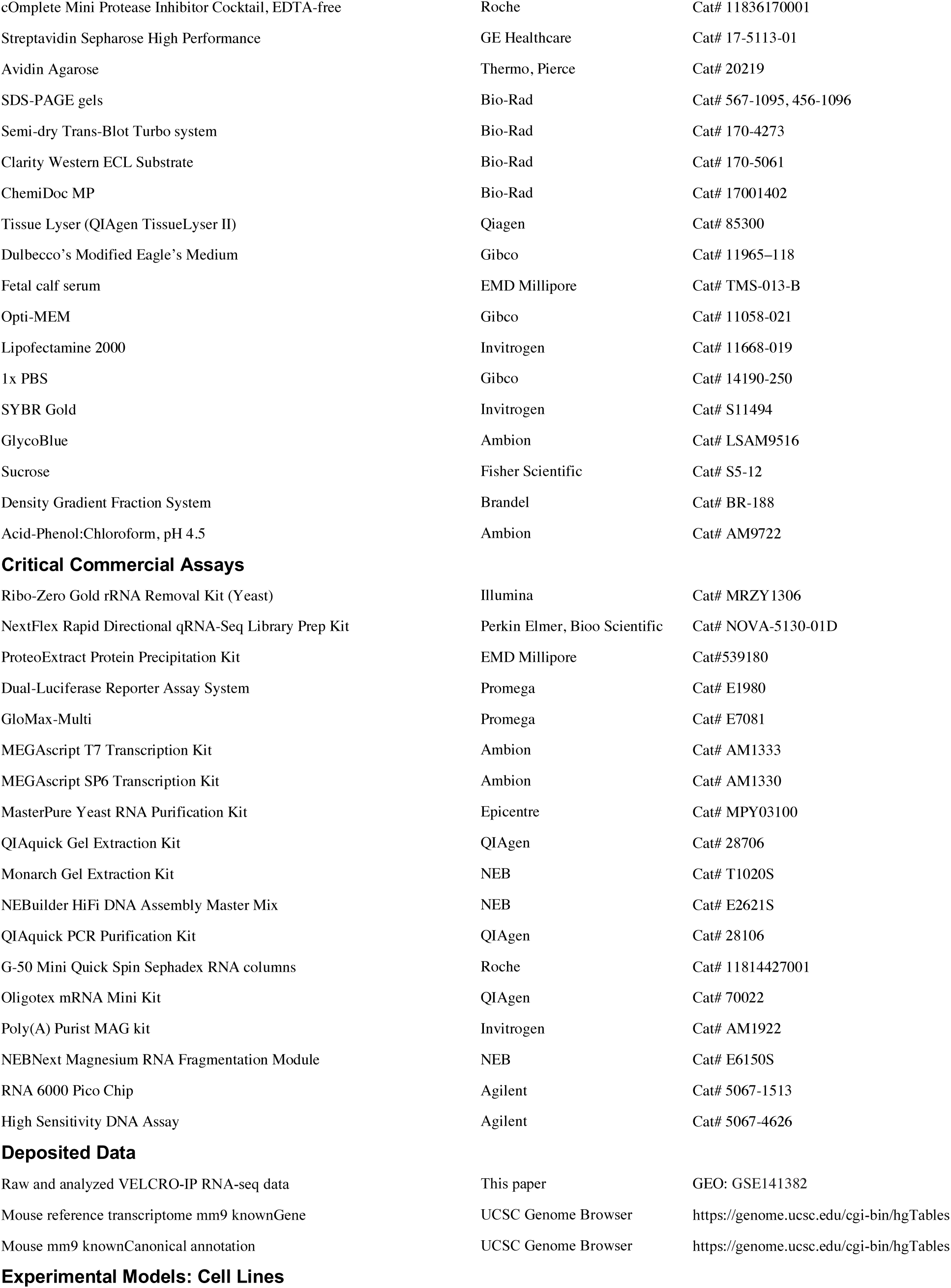
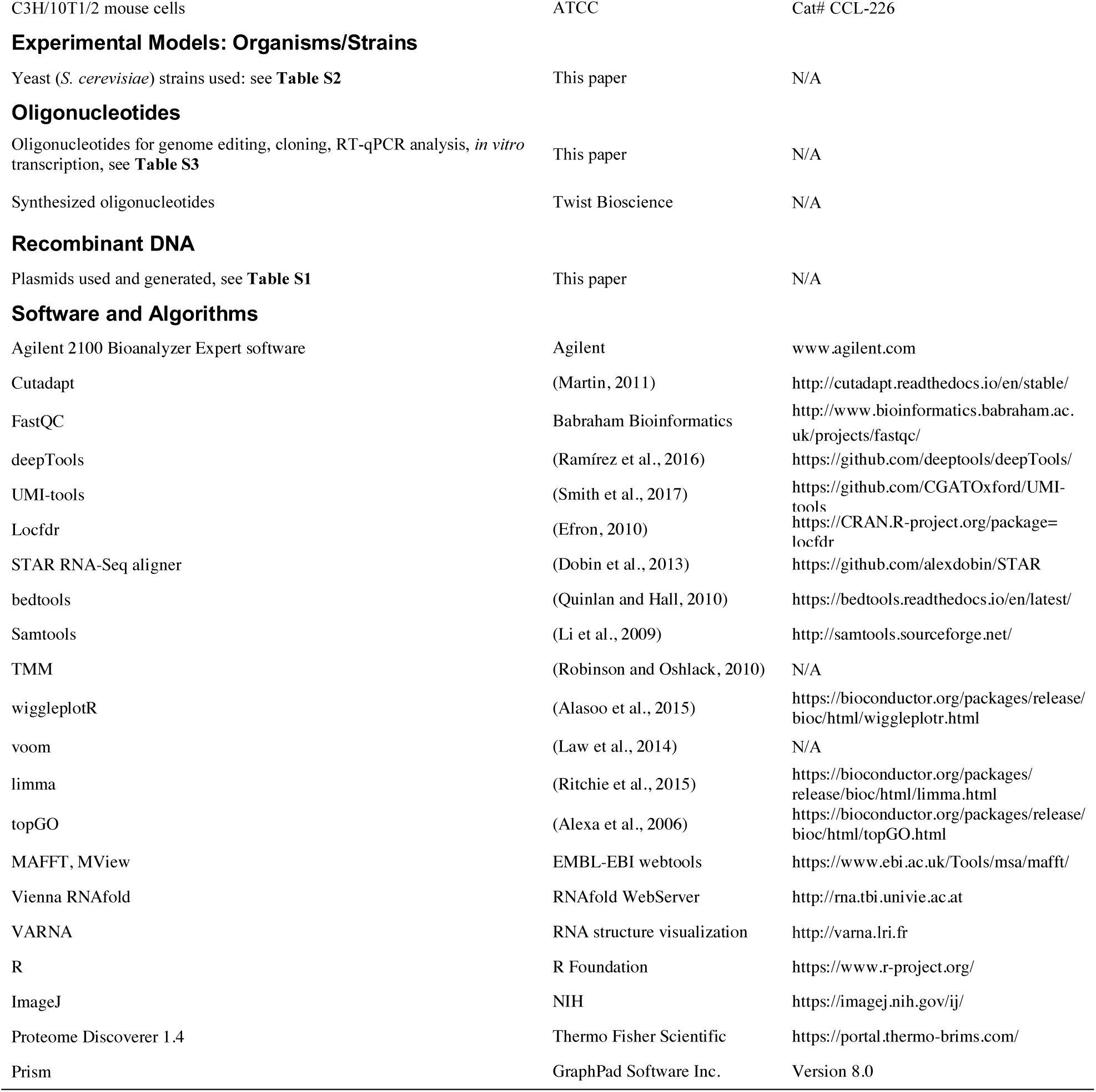

## SUPPLEMENTAL TABLES

**Table S1:**
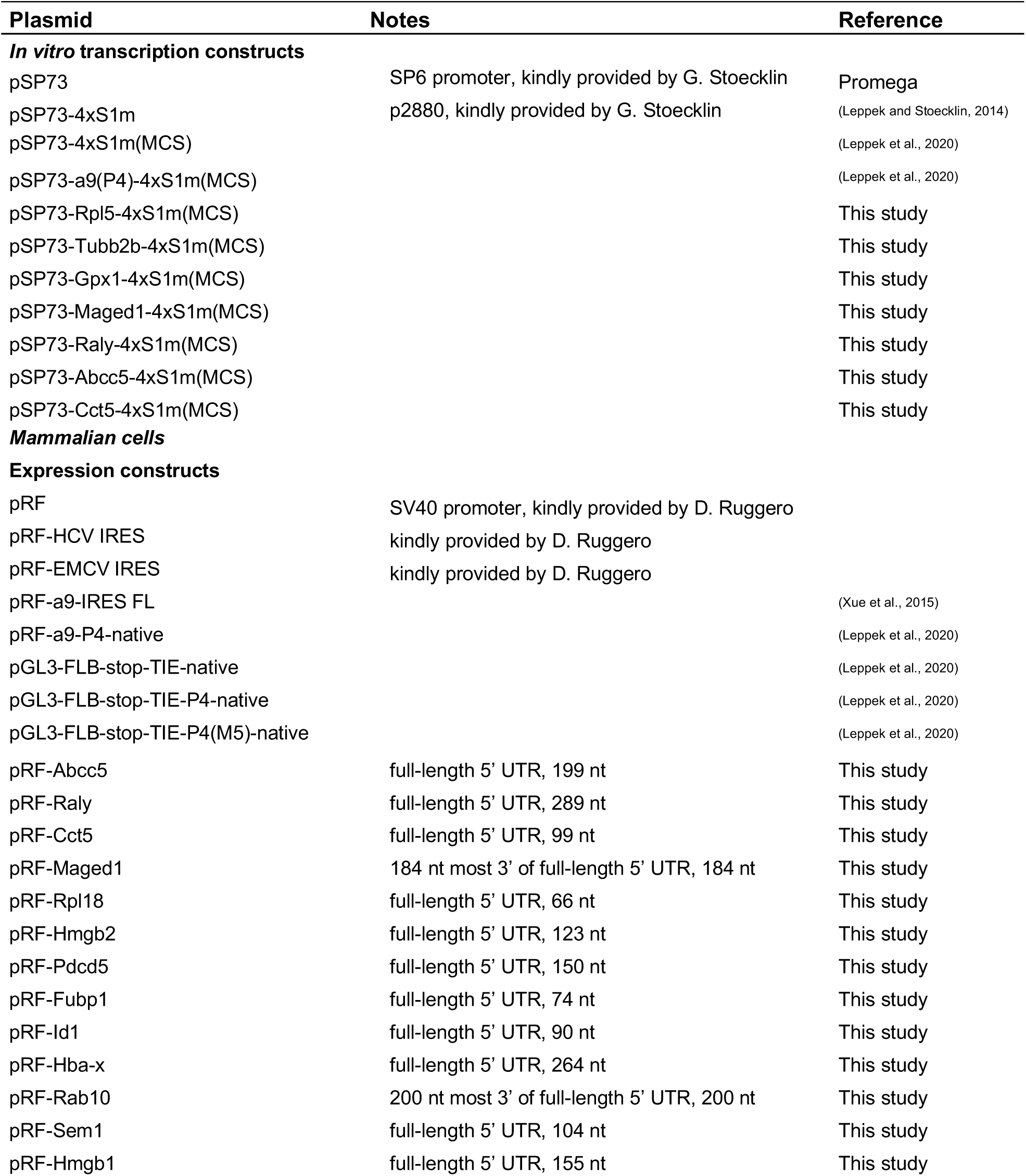
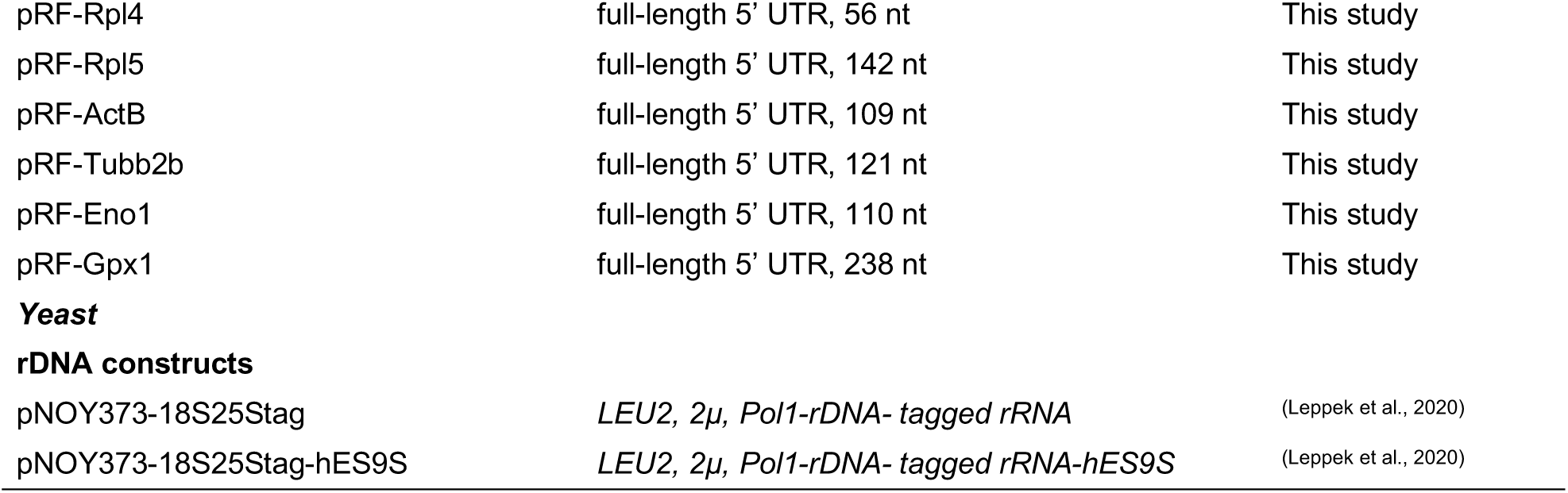
Plasmids used in this study All plasmids used for *in vitro* transcription and mammalian transient transfection or yeast transformation are listed in the table.

**Table S2:** Yeast strains used in this study All yeast strains used and/or generated for this study are listed in the table.

| Table S2. List of yeast strains used in this study |  |  |
| --- | --- | --- |
| Strain | Genotype and Notes | Reference |
| KAY488<br>(NOY890) | *MATA ura3-1 leu2-3,112 his3-11 trp1-1 ade2-1 can1-100 rdna$\Delta\Delta$::HIS3*<br>carrying *pRDN-hyg::URA3* | (Nemoto et al., 2010) |
| NOY890<br>WT rRNA | *MATA ura3-1 leu2-3,112 his3-11 trp1-1 ade2-1 can1-100 rdna$\Delta\Delta$::HIS3*<br>carrying *pNOY373-WT rRNA::LEU2* | (Leppek et al., 2020) |
| NOY890<br>tagged-hES9S | *MATA ura3-1 leu2-3,112 his3-11 trp1-1 ade2-1 can1-100 rdna$\Delta\Delta$::HIS3*<br>carrying tagged *pNOY373-rRNA-hES9S::LEU2* | (Leppek et al., 2020) |
| RPS2-Flag | *MATA ura3-1 leu2-3,112 his3-11 trp1-1 ade2-1 can1-100 rdna$\Delta\Delta$::HIS3*<br>*RPS2-Flag::kanMX6* carrying *pRDN-hyg::URA3* | This study |
| RPS2-Flag<br>WT rRNA | *MATA ura3-1 leu2-3,112 his3-11 trp1-1 ade2-1 can1-100 rdna$\Delta\Delta$::HIS3*<br>*RPS2-Flag::kanMX6* carrying *pNOY373-WT rRNA-hES9S::LEU2* | This study |
| RPS2-Flag<br>tagged-hES9S | *MATA ura3-1 leu2-3,112 his3-11 trp1-1 ade2-1 can1-100 rdna$\Delta\Delta$::HIS3*<br>*RPS2-Flag::kanMX6* carrying *pNOY373-tagged rRNA-hES9S::LEU2* | This study |

**Table S3.**
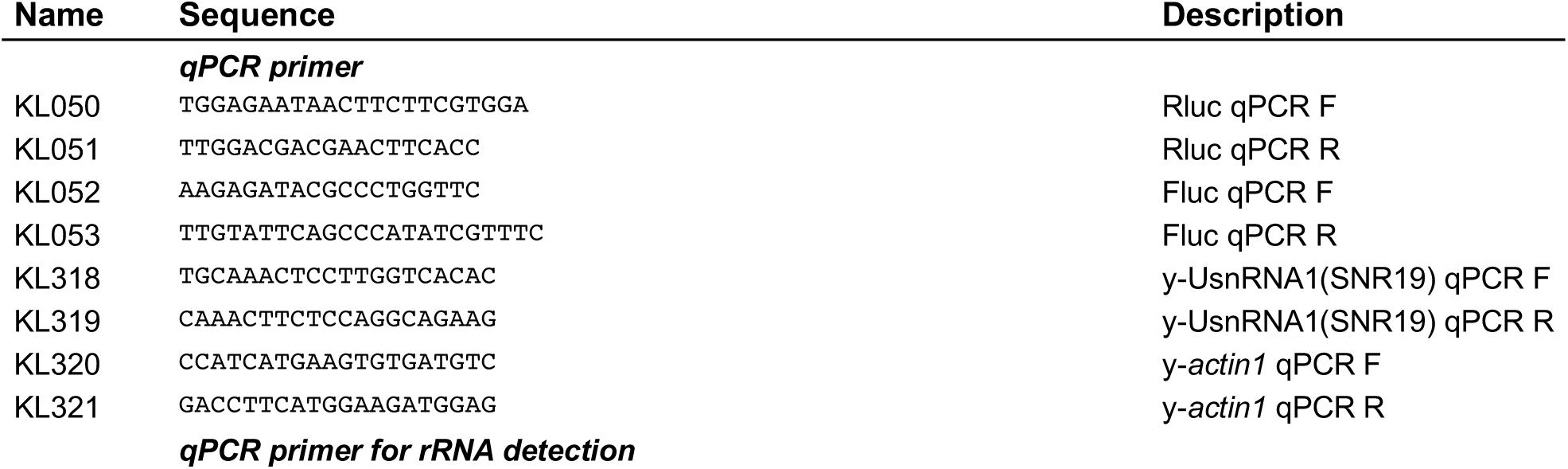
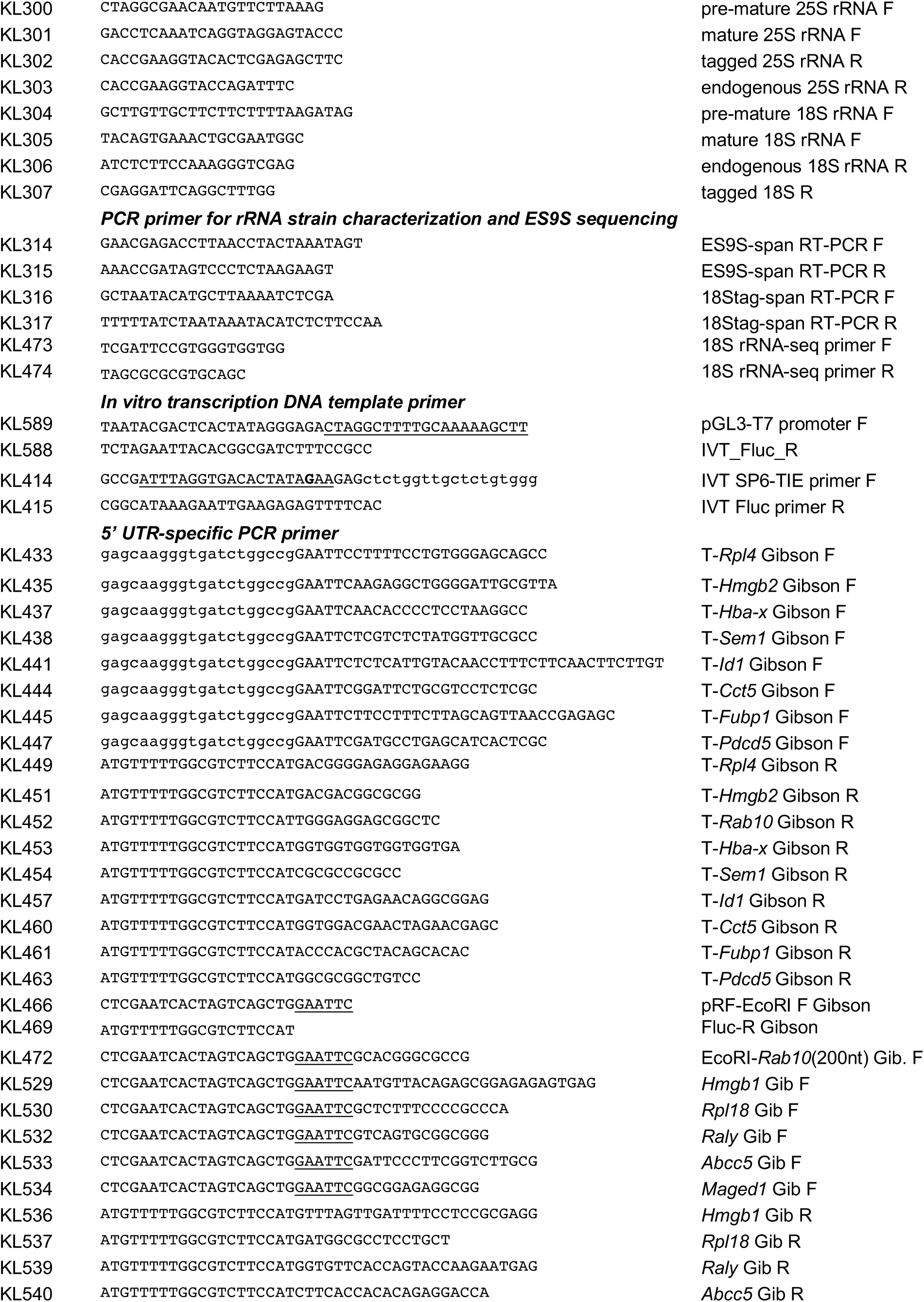
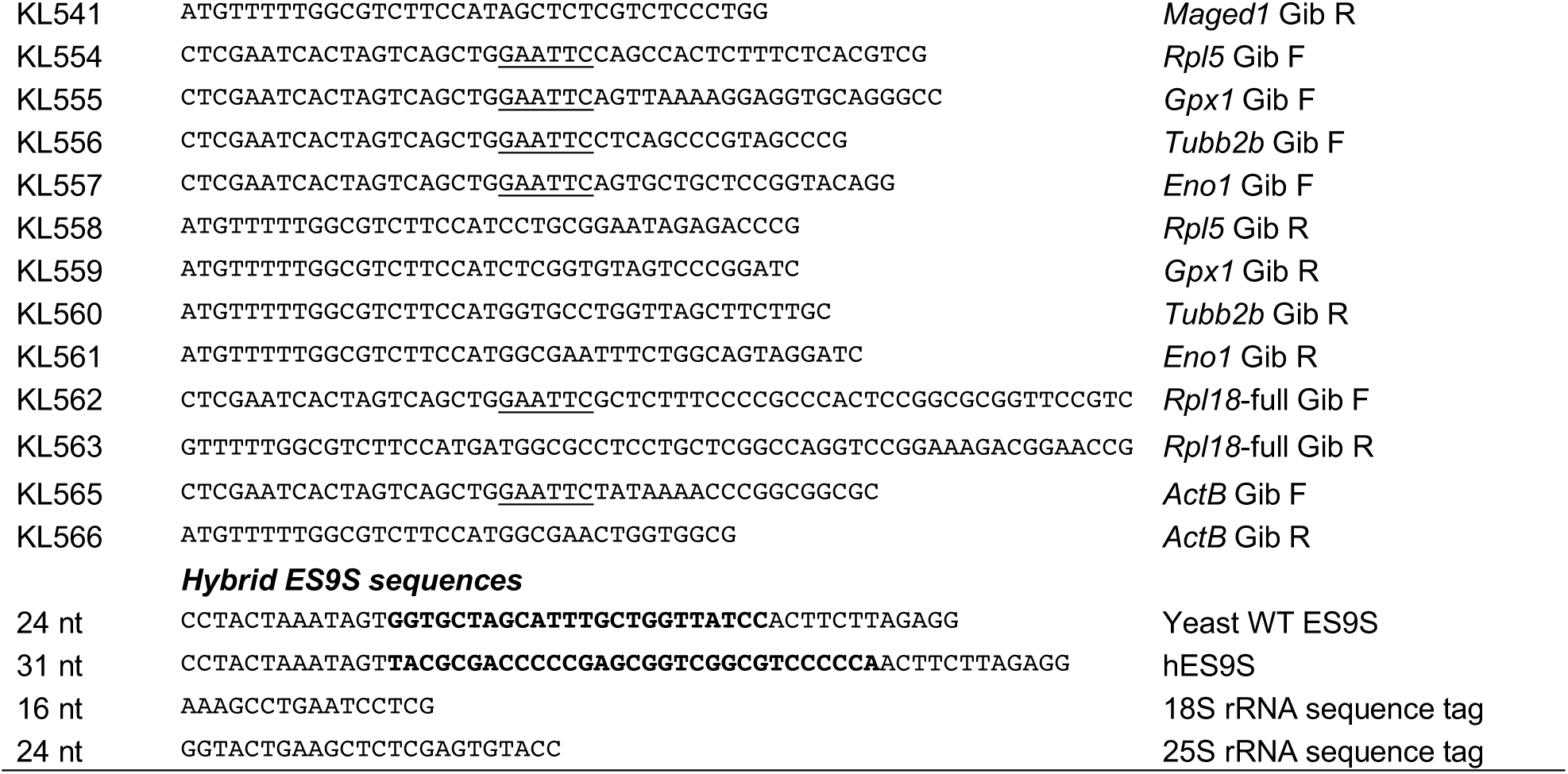
DNA Oligonucleotides used in this study All DNA oligonucleotides used for cloning, RT-PCR, and RT-qPCR are listed in the table. F, forward primer; R, reverse primer.

**Table S4. mRNA regions significantly enriched by VELCRO-IP RNA-seq** RNA was extracted after IP of yeast WT- and hES9S-ribosomes and subjected to RNA-seq in three biological replicates. The table shows all mRNA regions significantly enriched by hES9S-ribosome-IP. We referred to a cutoff of FDR<0.05 for selection of mRNA 5’ UTR targets for experimental validation and for analysis of enrichment of mRNA regions (**Figure 4E**) and of GO terms (**Figure 4G**).

**Table S5. GO term analysis for all genes enriched with hES9S.** Analysis of enriched genes in all GO term categories given in **Figure 4G** and **Figure S6** are described in the table.

